# Bronze Age *Yersinia pestis* genome from sheep sheds light on hosts and evolution of a prehistoric plague lineage

**DOI:** 10.1101/2025.02.07.637078

**Authors:** Ian Light-Maka, Taylor R. Hermes, Raffaela Angelina Bianco, Lena Semerau, Pavel Kosintsev, Valeriia Alekseeva, Donghee Kim, William P. Hanage, Alexander Herbig, Choongwon Jeong, Christina Warinner, Felix M. Key

## Abstract

Most human pathogens are of zoonotic origin. Many emerged during prehistory, coinciding with domestication providing more opportunities for spillover from original host species. However, we lack direct evidence linking past animal reservoirs and human infections. Here we present a *Yersinia pestis* genome recovered from a 3rd millennium BCE domesticated sheep from the Eurasian Steppe belonging to the Late Neolithic Bronze Age (LNBA) lineage, until now exclusively identified in ancient humans across Eurasia. We show that this ancient lineage underwent ancestral gene decay paralleling extant lineages, but evolved under distinct selective pressures contributing to its lack of geographic differentiation. We collect evidence supporting a scenario where the LNBA lineage, unable to efficiently transmit via fleas, spread from an unidentified reservoir to humans via sheep and likely other domesticates. Collectively, our results connect prehistoric livestock with infectious disease in humans and showcase the power of moving paleomicrobiology into the zooarcheological record.

## Introduction

Zoonoses – infectious diseases that can spread from a non-human host to humans – have been identified by the World Health Organization as among the greatest modern threats to human health ^1^. They have been a leading cause of death and disease throughout the human past, and the majority of known pathogens are believed to be zoonotic in origin ^2^. In the early Holocene, humans began transitioning from foraging subsistence to food production, which increased opportunities for zoonoses through larger and denser communities and closer contact with domesticated livestock. In particular, the domestication of sheep, goats, pigs, and cattle and their cohabitation with people have been hypothesized as drivers for the emergence of deadly human pathogens causing infectious diseases as varied as tuberculosis ^3^, salmonellosis ^4^, measles ^5^, and plague ^6^. Reconstructed ancient pathogen genomes from human skeletal remains have identified progenitors to human-adapted lineages and a rise in zoonotic pathogens linked to Eurasian Neolithization and Bronze Age population expansions ^4,7,8^. However, despite the relevance of animals for understanding the emergence, reservoirs, transmission dynamics, and impact of pathogens on human history, the zooarchaeological record remains largely unexplored for ancient pathogens.

Reconstructing pathogen genomes from ancient animals, however, is challenging because, unlike humans, animal remains primarily enter the archaeological record as food waste ^9^. As such, not all skeletal elements are expected to be recovered and many of them will have been exposed to high heat through cooking and to the elements prior to archaeological deposition ^10^. Moreover, people tend to avoid consuming visibly sick animals ^11^, and therefore faunal assemblages are likely biased towards healthy animals. Even when infected animals are consumed, a single animal may infect many people, and the probability of that specific animal being found and later studied may be low. All of these factors decrease the chances of recovering pathogen DNA from the zooarchaeological record. To date, only a few ancient pathogen genomes have been reported from faunal remains ^12–14^, and only a single genome older than a thousand years has been recovered, from the zoonotic pathogen *Brucella melitensis ^15^*.

*Yersinia pestis*, the zoonotic bacterial pathogen responsible for multiple devastating pandemics throughout human history ^16^, has been well studied using ancient DNA, with nearly 200 genomes reconstructed from human remains to date ^17,18^. Yet only a single partial *Y. pestis* genome is available from an ancient animal—a rat—dated to the medieval period ^12^. All extant *Y. pestis* strains trace their origins to a common ancestor four millennia ago in the Samara region along the middle Volga River ^19^, which subsequently diversified into multiple lineages (branches 0-4) ^20^. The primary reservoir of extant *Y. pestis* has been small mammals, primarily sylvatic rodents, from whom the pathogen is transmitted via fleas to other sylvatic rodents and sporadically to larger animals and humans, causing the highly fatal disease bubonic plague ^21^. Genomic comparison and experimental studies of flea colonization have identified a combination of gene acquisition, gene loss, and pseudogenization events occurring since the evolution of *Y. pestis* from its ancestor, *Y. pseudotuberculosis*, that enabled more efficient vector borne transmission of *Y. pestis* by fleas to a broader host range, and contributed to its high human mortality, numbering in the millions ^16,22–26^.

Separate from the flea-adapted form of *Y. pestis* responsible for bubonic plague outbreaks, a basal form of *Y. pestis* known as the Late Neolithic Bronze Age (LNBA) lineage has been so far exclusively identified in dozens of human archeological remains across Eurasia ^8,27,28^. This presumably extinct LNBA lineage was maintained as a single lineage for over 2,000 years (ca. ∼2900-500 BCE), while infecting diverse human populations from Western Europe to Mongolia during a period of heightened pastoralist mobility and interaction throughout the Eurasian steppes ^29,30^. Surprisingly, the LNBA lineage lacks the key genetic features shown to drive flea transmission ^27^, which has so far obscured its ancient host reservoir and transmission dynamics.

Here, we analyze sheep and cattle remains from the site of Arkaim, a fortified Bronze Age settlement associated with the Sintashta-Petrovka culture in the southern Urals region of the Eurasian steppe ^31,32^ and identify a *Y. pestis* infection in a domesticated sheep directly dated to 1935-1772 cal BCE. The reconstructed pathogen genome belongs to the LNBA lineage, and it phylogenetically clusters with contemporaneous genomes of the LNBA lineage isolated from human remains. We deduce its virulence from its ancestral gene content and test different transmission scenarios using phylodynamics and population genetic theory. In contrast to extant *Y. pestis*, we find that the LNBA lineage evolved under purifying selection but also exhibits signals of parallel evolution at its phylogenetic tips. Our results point to differences in how the LNBA lineage propagated within its reservoir and spread compared to extant *Y. pestis*, and we propose a model scenario that could explain its evolution and transmission in prehistoric human populations.

## Results

### *Y. pestis* DNA identified in a Bronze Age domesticated sheep

To explore the zoonotic range of *Y. pestis* during prehistory, we obtained and sequenced DNA from the skeletal remains of 23 domesticated sheep and cattle at the Middle Bronze Age site of Arkaim, Russia, and analyzed the resulting metagenomic datasets for the presence of *Y. pestis* DNA using HOPS ^33^ (**Figure 1A, Dataset S1**). In the dataset of sample ARK017, we identified a putative signal of ancient *Y. pestis* DNA (**Figure S1**). Next, we performed in-solution *Y. pestis* genome capture of six *Y. pestis* libraries produced from four independent DNA extractions from ARK017; each contained *Y. pestis* sequences (**Dataset S2**). Together, they yielded a 2.31-fold genome with 75.01% breadth based on alignment to the *Y. pestis* reference genome CO92 (2.26-fold chromosome, 3.86-fold pCD1, 2.64-fold pMT1, 10.37-fold pPCP1), which exhibited deamination patterns, an edit distance distribution, and short fragment lengths associated with a genuine ancient *Y. pestis* genome (**Figure S2**, **Table 1**).

**Figure 1.**
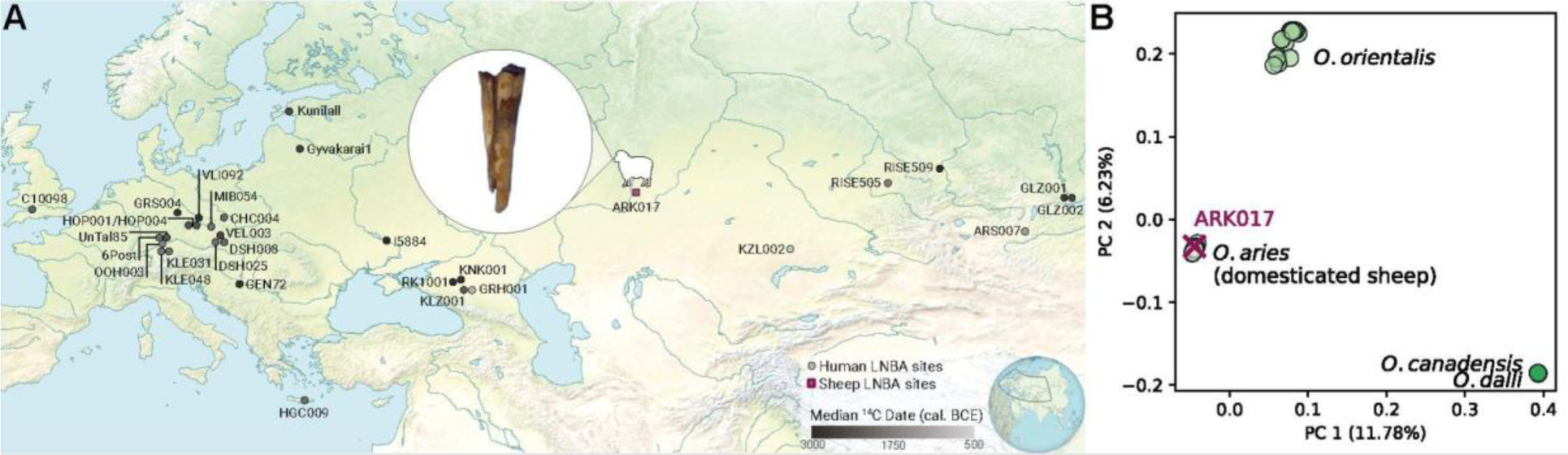
LNBA lineage *Y. pestis* identified among ancient domesticated sheep and humans. **A.** Recovered LNBA lineage *Y. pestis* genomes from a sheep (sheep icon, square) and previously published humans (circles) over a time transect of 2,500 years. Site color is scaled by median cal. BCE date. Circle inset shows sheep tooth specimen ARK017. **B.** ARK017 is genetically identified as a domesticated sheep (*Ovis aries)*, rather than a wild sheep, via projection of ARK017 into *Ovis* ntDNA principal components 1 and 2. SNVs were generated from the International Sheep Genomics Consortium reference panel of *O. aries* and wild sheep species. See also **Figure S3**.

**Table 1:** ARK017 *Y. pestis* genome reconstruction. Mapping statistics for merged data from in-solution *Y. pestis* DNA enrichment libraries and after applying mapping quality threshold of 30. See also **Figure S2** and **Dataset S2**.

| Sample Name | Deduplicated Mapped Reads | Mean coverage | Percent Breadth covered $\geq 1X$ | Percent Endogenous | Mean read length | Terminal 5' C to T misincorporation rate |
| --- | --- | --- | --- | --- | --- | --- |
| ARK017 | 544,282 | 2.31 | 70.50% | 0.06% | 46.38 | 15.63% |

The ARK017 *Y. pestis* genome originates from a tooth taxonomically assigned by zooarchaeological methods to *Ovis* (**Figure 1A, inset**). Because a secure taxonomic assignment for caprine ^34^ and bovine species ^35^ can be challenging from incomplete remains, we examined the mitochondrial DNA of the 23 animal samples and aligned them to a database of 34 mitochondrial genomes of wild and domesticated ungulates with habitat ranges in central Eurasia (also including human and dog) using MALT ^36^. Eleven samples were assigned to *Bos*, and 12 samples were assigned to *Ovis*, including ARK017 (**Dataset S1**). To further determine whether the ARK017 tooth originates from a domesticated sheep (*Ovis aries*) or a wild relative (e.g., *O. orientalis*, *O. dalli*, *O. canadensis*), we projected ARK017 into the top principal components (PCs) calculated from a panel of present-day wild and domesticated sheep genomes aligned to *O. aries* (**Dataset S3**). ARK017 clusters with *O. aries* in the first and second principal components, confirming its identification as a domesticated sheep (**Figure 1B**). The third principal component distinguishes geographically distinct subgroups of *O. aries*, and ARK017 clusters with southwest Asian *O. aries* (**Figure S3B**), as expected based on the known history of cultural transmission and genetic connections between Neolithic southwest Asian farmers and Eneolithic-Bronze Age steppe pastoralists ^37–39^. Similar to the *Y. pestis* aligned reads, the *Ovis* aligned reads showed molecular damage consistent with an ancient origin (**Figure S3C,D**). The ARK017 tooth produced a radiocarbon date of 1935 - 1772 cal. BCE (3532 ± 19 BP; Methods), placing it firmly within the Sintashta period (2040 - 1730 BCE) of the Sintashta-Petrovka cultural complex during the Middle-Late Bronze Age ^40–42^.

### Sheep *Y. pestis* genome is part of the LNBA lineage

To determine the phylogenetic placement of the prehistoric ARK017 *Y. pestis* genome, we compared it to 66 ancient *Y. pestis* genomes together with 134 modern genomes representing the known diversity of the species (**Dataset S4**). All genomes were aligned to the CO92 *Y. pestis* reference genome, and 4,444 high-quality SNVs were retained for phylogenetic analysis (Methods, **Dataset S5**). We constructed a Maximum Likelihood phylogeny, which placed the ARK017 genome within the LNBA lineage (**Figure 2A**). Focusing on the LNBA lineage, the sheep-derived ARK017 *Y. pestis* genome falls among contemporaneous human-derived genomes, with DSH025 (2026-1884 cal. BCE, Austria) falling basal and MIB054 (1952-1897 cal. BCE, Czech Republic) being more phylogenetically derived (**Figure 2B**). While the LNBA lineage phylogeny has on average a very high bootstrap support (≥99%), ARK017 has a bootstrap support of 78%; the alternative trees reveal ambiguous branching with neighboring, contemporaneous genomes DSH025 and KLE031 (2016-1891 cal. BCE, Germany). Interestingly, the phylogenetic placement and calibrated ^14^C date for ARK017 is consistent with the pattern of repeated genetic replacement within the LNBA lineage despite the large geographic range of contemporaneous infections. We note that the calibrated ^14^C dates of the six most basal LNBA lineage genomes (RISE509 through GRS004) largely overlap, suggesting those infections happened within a time window of approximately 200 years ranging from Central Asia to Western Europe (**Figure 1A**, **Figure 2B**). Similar to all previously reconstructed LNBA lineage genomes derived from human remains, ARK017 lacks the *ymt* gene necessary for efficient flea transmission ^24^, and it follows other previously identified pathogenicity gene presence-absence patterns consistent with its phylogenetic location within the LNBA lineage (**Figure S4**) ^8^. Hence, we provide the first LNBA lineage *Y. pestis* genome derived from a non-human host – a sheep, thus enabling the exploration of *Y. pestis* zoonoses and evolutionary dynamics across species.

**Figure 2.**
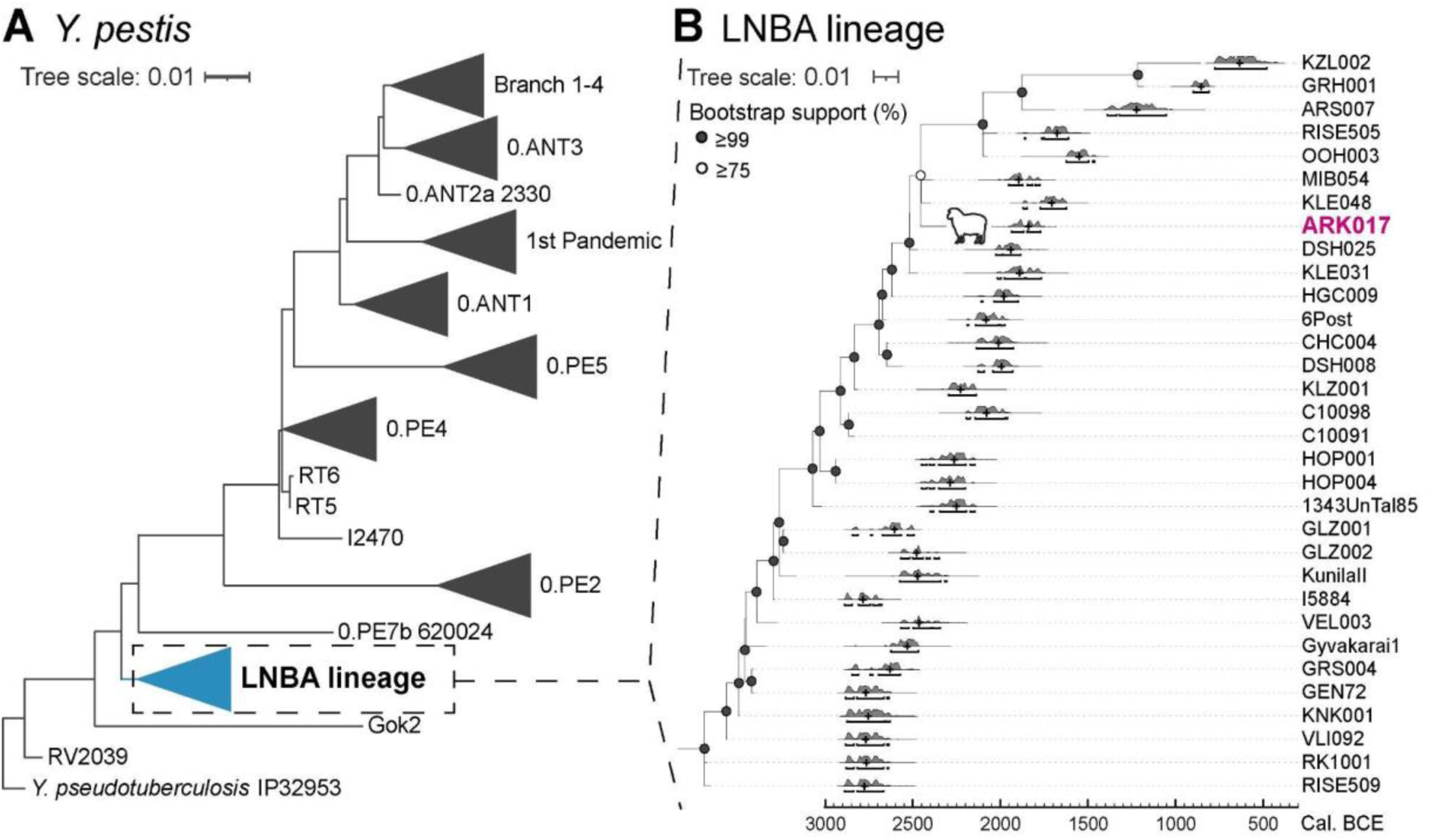
Recovered *Y. pestis* genome from ancient sheep, ARK017, confidently placed within the LNBA lineage diversity. **A.** Maximum likelihood *Y. pestis* tree generated using RAxML with a GTR+G model using 4,444 high quality variant positions across 200 *Y. pestis* genomes from the ancient and extant diversity, rooted by the *Y. pseudotu*berculosis isolate IP32953. Collapsed Branch 1-4 also contains 2nd Pandemic genomes. Genomes included in 1st and 2nd Pandemic branches can be found in **Dataset S4**. **B.** The ARK017 Y. pestis genome (purple label, sheep icon) is placed within the LNBA lineage diversity (blue collapsed branch in panel A), with 78% bootstrap support, over 10,000 replicates. Bootstrap support values above 75% are marked on internal tree nodes with an empty circle, and gray circles represent bootstrap support ≥ 99%. ^14^C sample ages for LNBA lineage genomes are plotted as a probability distribution over calibrated Before Common Era (cal. BCE) date ranges using OxCal v.4.4 ^43^ with the IntCal20 atmospheric curve ^44^. Regions making up 95.4% date range distribution are marked below in bars, and the median date estimate is marked with a vertical tick. Collapsed trees were generated using Interactive Tree of Life ^45^. For an uncollapsed tree, see data availability statement in methods.

### Parallel genomic decay of LNBA lineage *Y. pestis* but retention of genes impacting virulence in its ancestor

The genetic repertoire of *Y. pestis* is known to differ significantly from its less virulent ancestor, the enteric pathogen *Y. pseudotuberculosis*, with differences driven primarily through genomic decay ^22,46^. Genomic decay (pseudogenization or loss of genes) has been linked to *Y. pestis* transmission potential within the flea vector, its virulence, and improved fitness for its intracellular life cycle ^24,47–49^. The LNBA lineage represents an early diversifying clade in the *Y. pestis* phylogeny. Its genetic origin is estimated to be separated by approximately 500 years from the inferred most recent common ancestor of all known *Y. pestis*, which in turn presumably split from *Y. pseudotuberculosis* more than forty thousand years ago ^27^. Ancient DNA allows us to timestamp the evolutionary events from the *Y. pseudotuberculosis* ancestor to the highly virulent *Y. pestis* known today, which in turn helps us to better understand the virulence of the LNBA lineage, which is believed to have gone extinct more than 2,000 years ago.

To investigate the timing and extent of ancestral gene content retention, we identified genes relevant for *Y. pseudotuberculosis* biology by characterizing its core genome (3081 genes) and assessed their presence and loss across different clades of *Y. pestis*. Interestingly, the LNBA lineage and other basal ancient genomes show a greater affinity with *Y. pseudotuberculosis* than any extant *Y. pestis* clade in a PCA based on normalized gene breadth (**Figure 3A**), suggesting a retention of ancestral genes that agrees with a recent analysis focusing instead on the *Y. pseudotuberculosis* complex pangenome ^50^. However, it is possible that inadvertent cross-mapping from orthologous sequences present in the ancient metagenomic datasets could bias such analyses. To explicitly check for this bias, we estimated retention of *Y. pseudotuberculosis* core genes across different ancient datasets grouped by time. We observe that other prehistoric *Y. pestis* genomes contemporaneous with the LNBA lineage do not differ from the LNBA lineage in core gene content (p-value = 0.754, Mann-Whitney-U test); in contrast, both the ancient genomes from the historical period (First and Second Pandemics, phylogenetically related to Branches 0-4) and the extant *Y. pestis* diversity retained significantly fewer *Y. pseudotuberculosis* core genes compared to the LNBA lineage (both p-value < 0.005, Mann–Whitney-U test; **Figure 3B**). In addition, we observe a time-dependent degradation of *Y. pseudotuberculosis* gene breadth on the LNBA lineage (r^2^=0.83), which would be unexpected if driven by random cross-mapping and suggests a clock-like loss of ancestral genes. These findings are in agreement with the separation of the prehistoric and historic ancient *Y. pestis* genomes in the PCA and strongly suggest that the most ancient *Y. pestis* genomes indeed retained a larger repertoire of *Y. pseudotuberculosis* genes than is now represented among extant lineages of *Y. pestis*. The time dependence of genome decay is noteworthy, as it parallels previous observations of pseudogenization through time on the LNBA lineage ^8^, and reveals that the process of *Y. pseudotuberculosis* genome decay was dynamic across early diversifying ancient lineages.

**Figure 3.**
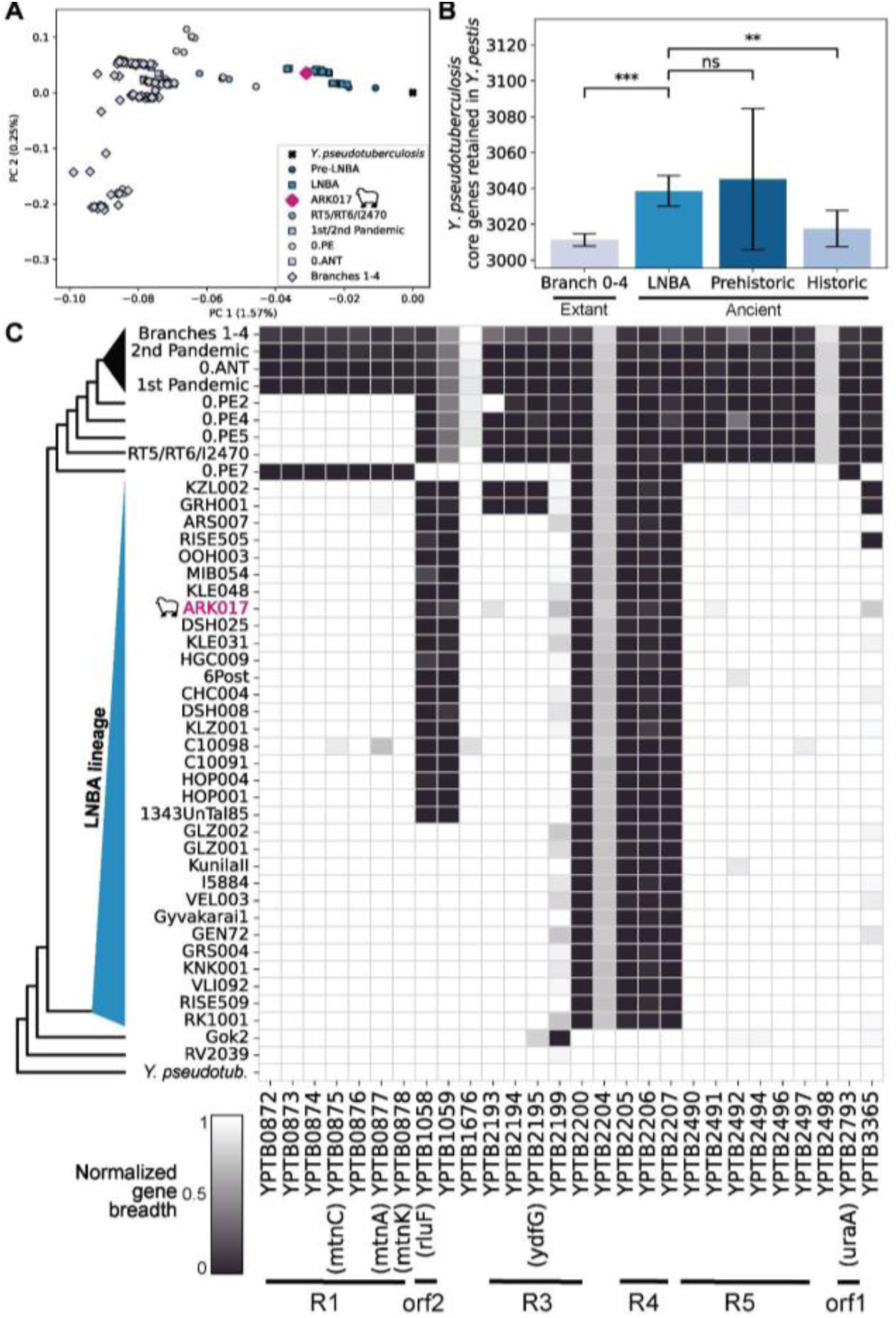
LNBA lineage retains unique gene configuration from its *Y. pseudotuberculosis* ancestor and underwent gene loss parallel to extant diversity during its evolution. A. LNBA lineage and basal samples associate more closely with the *Y. pseudotuberculosis* (IP32953) ancestor than any extant or more recent ancient genomes in a PCA based on gene breadth of core genes in *Y. pseudotuberculosis*. **B.** The LNBA lineage retained significantly more ancestral *Y. pseudotuberculosis* core genes than extant *Y. pestis* genomes and historical (1st/2nd pandemic) ancient genomes. No difference is observed when comparing the LNBA lineage to other prehistoric ancient genomes (contemporaneous or basal to the LNBA lineage), confirming a true retention of ancestral *Y. pseudotuberculosis* core gene content instead of orthologous sequences present in ancient metagenomic datasets. *** represents p-values < 0.0005, calculated using a Mann-Whitney-U test. Bars represent 95% confidence intervals of the mean across samples. Samples with strict-core gene presence estimates below two standard deviations of all samples were excluded (Methods). C. *Y. pseudotuberculosis* genic regions experienced repeated, independent genomic decay during early *Y. pestis* differentiation. Genes were identified within the *Y. pseudotuberculosis* core genome when present (breadth >99%) in at least one LNBA lineage or basal (RV2039, Gok2) genome and absent (breadth <90%) in at least 75% of other genomes. Genomes are phylogenetically ordered, simplified on the left for visual clarity, where *Y. pseudotub.* is the *Y. pseudotuberculosis* (IP32539) outgroup. For each collapsed branch, the maximum observed breadth measurement across all genomes per row is plotted. Core genes belonging to previously described regions are marked at bottom with bars, some of which impact *Y. pseudotuberculosis* virulence and growth rate *in vivo ^46^*, full regions can be found in **Figure S5, Table S1.** Agreement of ARK017 with placement in LNBA lineage also seen in *Y. pestis* virulence genes **(Figure S4)**. For B and C, sample group information can be found in **Dataset S4**.

We next sought to characterize the individual *Y. pseudotuberculosis* core genes retained in the *Y. pestis* basal ancient diversity and the LNBA lineage when compared to the extant diversity. We filtered for *Y. pseudotuberculosis* core genes that were present (breadth > 99%) in at least one LNBA lineage or basal ancient sample and absent or likely pseudogenized (breadth < 90%) in almost all extant *Y. pestis* (> 75% samples). We identified 28 genes (**Figure 3C**), some of which were previously thought to be completely absent in *Y. pestis ^22,46^*. We find evidence for repeated and independent (i.e., parallel) gene deletions in *Y. pestis* for half of the genes (n=14), six of which were independently lost within both the LNBA lineage and in early contemporaneous forms of flea-adapted *Y. pestis* (**Figure 3C**). Such parallel gene loss may be linked to shared evolutionary pressure for genomic decay in pathogens ^51^. Interestingly, two genomic regions in which we observe parallel gene loss, known as R3 and *orf2* ^22^, result in growth inhibition and attenuated virulence when deleted within the *Y. pseudotuberculosis* ancestor ^49^(**Figure 3C**). Notably, we observe stepwise deletion of multiple genes in the R3 region within the LNBA lineage, meaning that the region was not lost as a single genomic segment during *Y. pestis* evolution, but rather underwent repeated gene loss, paralleling the stepwise loss of the region during extant *Y. pestis* differentiation ^49^.

Lastly, the sheep-derived ARK017 genome shares a similar genetic make-up with contemporaneous human LNBA lineage genomes. This holds true for both the pattern of genomic decay (**Figure 3C**) and the *Y. pestis*-specific acquisition of genes associated with virulence and flea-borne transmission (**Figure S4**) ^8,27^. Taken together, these changes indicate a departure of the LNBA lineage in terms of virulence from its *Y. pseudotuberculosis* ancestor, as well as a deviation from the virulence of later *Y. pestis*. This raises the question of whether the life cycle and host ranges of the LNBA lineage also differed from modern *Y. pestis*, emphasizing the importance of further study of the roles of these *Y. pseudotuberculosis* genes in *Y. pestis* infection and ecology.

### Sheep and human LNBA lineage infections arose from the same source population

Despite the availability of more than 30 human-derived LNBA lineage *Y. pestis* genomes, the transmission dynamics leading to human infections has remained enigmatic ^8^. The reconstructed *Y. pestis* genome from a domesticated sheep in this study provides a novel opportunity to understand whether livestock contributed to the transmission of the LNBA lineage. Human-derived *Y. pestis* genomes of the LNBA lineage show a strong correlation between temporal and genetic distance (r^2^ = 0.93; shown in blue in **Figure 4A**), as has been previously shown ^8^ A strikingly similarly correlation is found in pairwise comparisons between the sheep infection, ARK017, and human infections (r^2^ = 0.91, shown in purple in **Figure 4A**), with the sheep-human pairwise comparisons of genetic distance placing it well within expectations based on human-human pairwise comparisons (p-value = 0.531, Mann-Whitney-U test; **Figure S6**). Moreover, ARK017 is most closely related to contemporaneous ancient human-derived genomes on the LNBA lineage phylogeny (**Figure 2B**), and its gene content is consistent with this placement (**Figure 3C, Figure S4**). All together, the commonalities in the molecular clock, phylogenetic placement, and gene content strongly support that both human and sheep infections arose from the same source population of the LNBA lineage, and may indicate that animal husbandry elevated the possibilities for transmission into humans.

**Figure 4.**
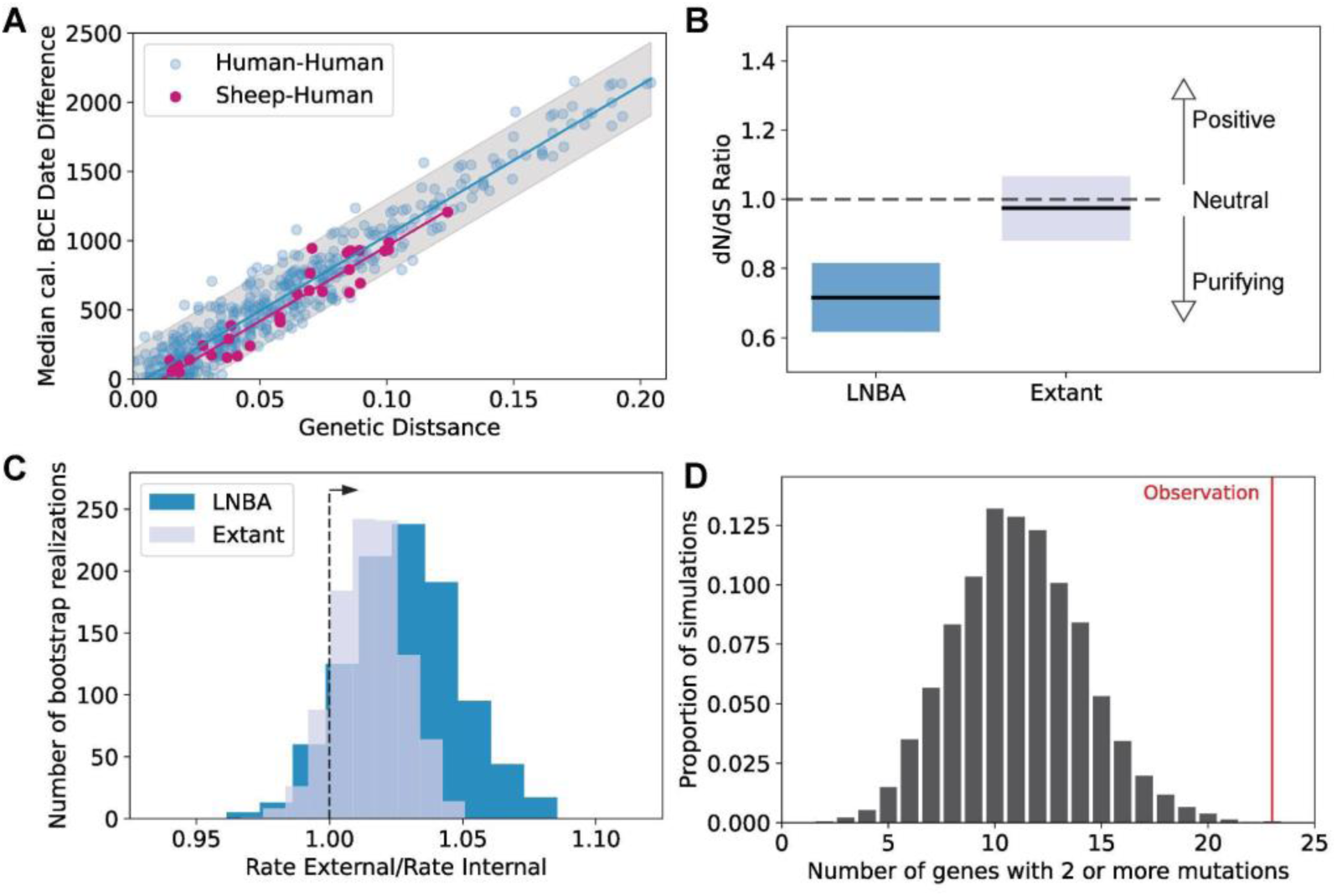
Domesticated sheep and human infections arose from a single pathogen source population, with the LNBA lineage evolving under purifying selection but with relaxed constraint during spillovers. **A.** Human infections (blue) of the LNBA lineage form a single lineage with a strong correlation (r^2^ = 0.93) between pairwise genetic distance (branch length) and temporal distance (median cal. BCE). Pairwise comparisons between sheep (ARK017) and human infections (purple) are likewise highly correlated (r^2^ = 0.91), indicating that they are sampled from the same source population. Regression line of human-human comparisons is shown in blue, with two standard deviations of the residuals shaded (gray); regression line of sheep-human comparisons is shown in purple. See also **Figure S6**. **B.** dN/dS measurements reveal signatures of purifying selection on the LNBA lineage, with an excess of silent, synonymous mutations, in contrast to extant lineages in which observed nucleotide diversity is consistent with neutrality. dN/dS values were calculated on all SNVs accumulated compared to their inferred ancestral state at the MRCA of the extant branches and LNBA lineage. **C.** Substitution rate of nonsynonymous external branch mutations (singletons - potential dead end mutations following spillover) compared to nonsynonymous internal branch mutations (shared - acquired during maintenance within the reservoir). The distribution represents 1,000 resamplings with replacement. Ratios above 1 (dashed arrow) indicate increased rates of nonsynonymous mutation accumulation on external branches. Non-significant elevated rate is identified for both the LNBA lineage and extant lineages, pointing at relaxed evolutionary constraint. **D.** Nonsynonymous mutations on external branches of the LNBA lineage (254 mutations) mutate 23 genes multiple times (red line). Compared to 10,000 simulations of the same number of mutations randomly distributed across all genic regions, the LNBA lineage nonsynonymous mutations hit the same genes at higher rates than expected by chance (p-value < 0.001). Observation remains significant when excluding the sheep infection ARK017 or KZL002, the youngest recovered genome contributing 36% of all singletons.

### Natural selection differentiates the LNBA lineage and extant lineages

Natural selection exerted by host immunity, transmission bottlenecks, and epidemic dynamics shape the genetic variation and genealogies of pathogens ^52^. Despite the vast spatial spread and large temporal range of all known LNBA lineage genomes, we do not observe any phylogeographic structure but instead a ‘ladder-like’ single-lineage genealogy ^8^. This is a hallmark of microbial pathogens that experience repeated replacement of existing diversity, which can be caused through bottlenecks (contraction of the population size), natural selection on the pathogen leading to the loss of previous diversity as it is outcompeted (positive selection) or unfit (purifying selection), or a combination thereof ^52,53^. We searched for signals of natural selection acting on the LNBA lineage, using the canonical measure of selection, the dN/dS ratio. The dN/dS ratio measures the rate of nonsynonymous to synonymous amino acid changes and has an expected value of 1 under neutrality, whereas less or more than 1 indicate purifying or positive selection, respectively. We identify an excess of silent, synonymous mutations on the LNBA lineage (dN/dS = 0.71, 95% confidence interval: 0.62 - 0.82; **Figure 4B**) compatible with purifying selection instead of positive selection. The dN/dS of extant *Y. pestis*, in contrast, is indistinguishable from neutrality (dN/dS = 0.97, 95% confidence interval: 0.88-1.07; **Figure 4B**), which is consistent with its diversification into multiple lineages (branches 0-4) ^20,54^. The lack of overlap between the two dN/dS estimates strongly suggests that, compared to the extant diversity of *Y. pestis*, the LNBA lineage experienced greater evolutionary pressure removing deleterious, amino acid changing mutations, independent of whether or not population bottlenecks also played a role in the phylogenetic patterns. Notably, the evolution of LNBA lineage *Y. pestis* thereby differs from that of other well-known pathogens who share a ladder-like phylogeny, including seasonal influenza, whose evolution is characterized by adaptive fixation of mutations over successive years that enable evasion of existing immunity ^55^.

While a mix of evolutionary pressure and possible bottlenecks on the LNBA lineage may explain the ladder-like shape of the reconstructed genealogy, it does not further our understanding of the transmission modes of the LNBA lineage. Today, *ymt*-positive *Y. pestis* is maintained in an enzootic-epizootic cycle, alternating between a latent enzootic stage in the rodent reservoir and an epizootic stage with rodent-flea *Y. pestis* exchange associated with high bacteremia in rodents that in turn facilitates flea-borne transmission into humans or other mammals ^56,57^. Explosive growth of a pathogen population during an epidemic leads to a fast but transient accumulation of non-synonymous, deleterious mutations due to incomplete purifying selection ^53,58–60^, which was previously proposed to underlie observed variation in the molecular clock between extant branches of *Y. pestis* ^20^. If the *ymt*-negative LNBA lineage was maintained in a comparable alternating cycle where the reconstructed genomes stem from dead-end infections during epizootic outbursts, we would expect more nonsynonymous, potentially deleterious mutations on external branches (dead-ends) compared to internal branches (successfully inherited). We investigated the ratio of rates of nonsynonymous mutations between external and internal nodes on both the LNBA and extant lineages, and we indeed observe an elevation, though non-significant, of nonsynonymous mutations on external lineages (ratio > 1) (**Figure 4C)**. This suggests that for both the LNBA lineage and extant diversity, sampling occurred during epizootic outbursts. Such an interpretation is consistent with observed long gaps between successful isolations of *Y. pestis* from extant plague foci, reservoir isolations clustering with human infections, and failures to isolate *Y. pestis* from live members of the reservoir population today ^61,62^.

Although the increased frequency of external, protein-altering mutations in the epizootic phase may be primarily deleterious, it is possible that some of those mutations provide an adaptive edge supporting the immune evasion, transmission, or disease severity in an incidental host while not competitive within the reservoir maintaining *Y. pestis*. Such a scenario, where the reconstructed infections encountered distinct evolutionary pressures than in the reservoir hosts, may be expected to produce signals of adaptive evolution hallmarked by genes carrying repeated, independent mutations ^63,64^. Indeed we observe an increase of genes with multiple, protein-altering external mutations compared to neutral simulations within the LNBA lineage (p-value<0.001, simulations; **Figure 4D**), a signature absent among extant human *Y. pestis* infections (p-value = 0.37, simulations) (**Figure S7A**). In contrast, a similar signature is absent among the internal, non-synonymous SNVs of the LNBA lineage (p-value = 0.313, simulations; **Figure S7B**), supporting the interpretation that different evolutionary pressures existed among the human and sheep infections compared to the reservoir. A possible adaptive role during spillover for the genes carrying repeatedly dead-end mutations can be deduced from their function, including known virulence factors upregulated during infection (*yapF* & *yapH*) ^65^, interference with inflammatory response (*yopM*) ^66^, a putative membrane protein (YPO2004) upregulated at human body temperature in *Y. pseudotuberculosis in vitro* ^67^, and a siderophore biosynthesis protein (YPO0776) previously identified to be evolving under diversifying selection across extant *Y. pestis ^68^* (**Table S2**). Importantly, the sheep infection ARK017 also carried nonsynonymous mutations in two of the genes with multiple mutations. This finding suggests that the sheep infection resulted in a comparable set of evolutionary constraints to the human infections and supports an interpretation where neither humans nor sheep were part of the LNBA lineage reservoir.

Taken together, we provide multiple lines of genetic evidence that the LNBA lineage experienced more purifying selection than extant *Y. pestis* lineages and that the reconstructed ancient human infections and ARK017 domesticated sheep infection were most likely a result of repeated epizootic outbursts representing evolutionary dead ends.

## Discussion

The identification of a Bronze Age *Y. pestis* genome from a domesticated sheep offers a novel perspective on the hidden evolution and host range of a prehistoric pathogen, and sets a precedent for the exploration of ancient diseases beyond humans. We show the sheep *Y. pestis* genome is part of the same single LNBA lineage, which lacks key genomic changes associated with efficient flea-borne transmission ^23,24,27^ and has until now only been reconstructed from prehistoric humans distributed across Eurasia^23,24,27^ Altogether, these factors raise questions about its transmission and maintenance as a single lineage across such a vast geographic space.

The domesticated sheep in this study comes from the Middle Bronze Age site of Arkaim, an important fortified settlement of the Sintashta-Petrovka cultural complex (ca. 2000–1600 BCE) in the southern Ural region of Russia. Pastoralism was the primary subsistence strategy of Sintashta-Petrovka sites and the economy at Arkaim centered on the husbandry of domesticated cattle, sheep, and horses ^69^. The Sintashta-Petrovka cultural complex was an early innovator of utilizing newly domesticated horses for battle, traction, and transport ^70^, and the use of horses in herding would have enabled pastoralists to cover a larger range and increase their livestock herd size up to 10-fold ^71^. These larger and more mobile herds likely brought them into greater contact with the wild fauna of grasslands, including hares and rodents, migratory birds, bovids (e.g., saiga antelope) and cervids (e.g., roe deer, elk) ^72^. Notably, the absence of cultivated crops or cereals at Sintashta-Petrovka sites ^73–75^ means that the inhabitants lacked the grain stores that are known to have attracted the kinds of commensal rodents later implicated in historic outbreaks of flea-adapted bubonic plague. Nevertheless, inhabitants would have frequently come into contact with wild rodents, such as marmots, while herding livestock or hunting rodents for their pelts and meat. In Central and East Asia today, local outbreaks of plague are frequently the outcome of skinning or eating infected wild rodents ^76,77^. Interestingly, grazing sheep in western China are also known to become infected with *Y. pestis* through licking or eating the carcasses of infected marmots, and in plague endemic regions up to 5% of sheep are seropositive for *Y. pestis* ^78^. Although often undetected, sheep plagues can spark local human outbreaks if the infected sheep are handled or consumed ^77,79^.

Given this epidemiology of plague among pastoralists in Asia today, in which sheep can be considered bridge hosts in transmission to humans ^80^, it is plausible that similar processes may have taken place in the propagation of the LNBA lineage, in particular among populations with well-developed animal husbandry (**Figure 5**). As such, flea transmission between the reservoir and humans would not have been necessary, but rather *Y. pestis* could have been transmitted from a wild host to humans through close contact with livestock or other domesticated animals. This would have limited the scale of local human outbreaks, and is consistent with the observation that LNBA lineage *Y. pestis* cases are associated with typical funerary deposition rather than catastrophic mass graves. While the ultimate reservoir of these infections remains unknown, distinct transmission routes in different species would be expected to lead to distinct selective pressures that could be reflected in *Y. pestis* genomes in agreement with the increased frequency of amino acid-changing mutations and the signature of parallel evolution identified among reconstructed infections.

**Figure 5.**
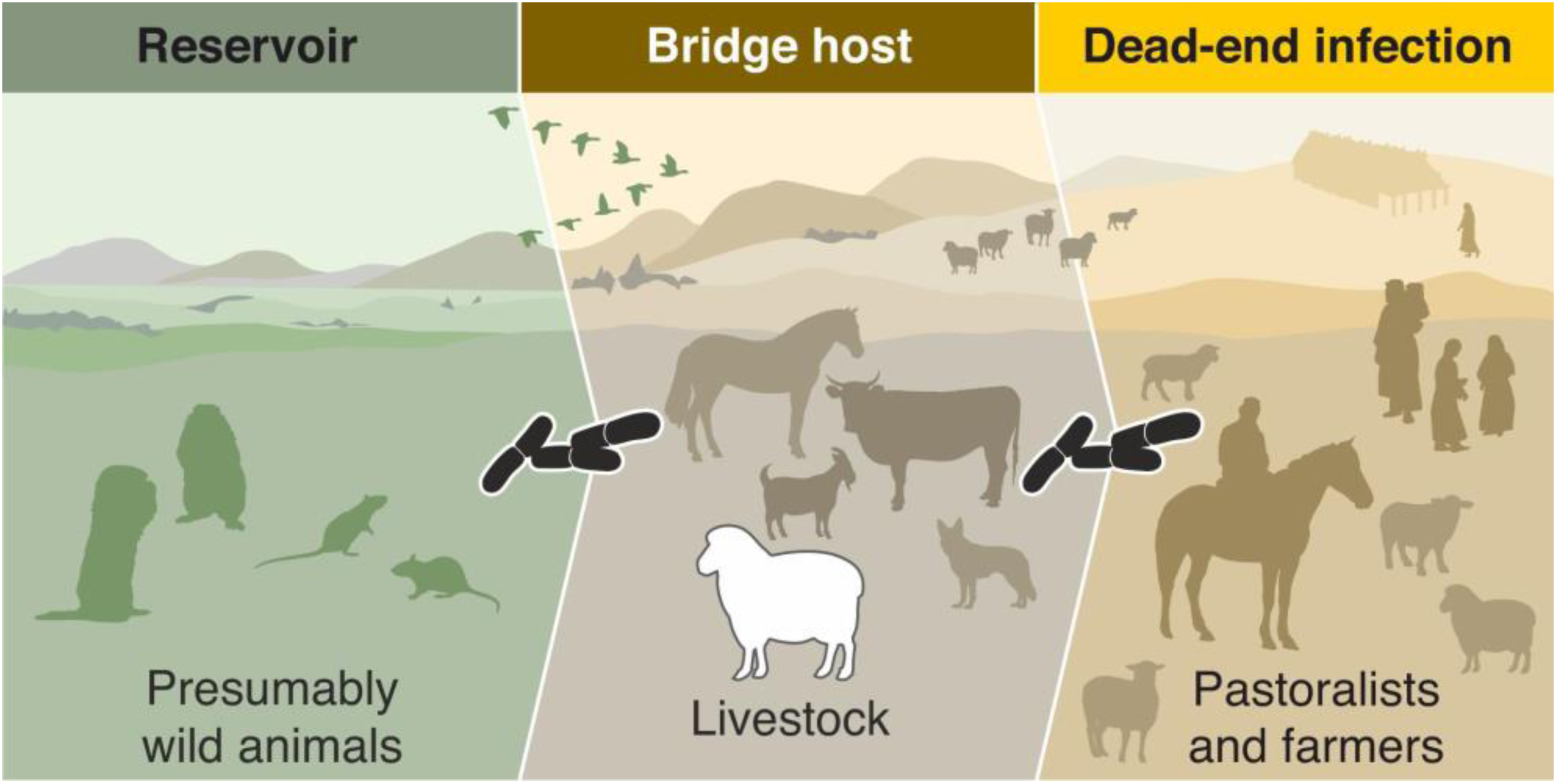
Conceptual scenario of the LNBA lineage transmission concluded from sheep and human derived *Y. pestis* genomes. Maintained among locally restricted reservoir populations, *Y. pestis* (shown in black) dispersed over large distances potentially through avian hosts. Domesticated animals, infected from the dispersal species or reservoir population, served as bridge hosts for dead-end infections in prehistoric human populations across Eurasia.

But how was the LNBA lineage maintained as a single lineage over thousands of years while spreading across Eurasia? The signature of purifying selection observed here provides an explanation for the ladder-like genealogy, but selective pressures alone do not shape phylogenetic patterns in natural populations ^52,53^. The influence of transmission bottlenecks during the enzootic-epizootic life cycle on the genetic diversity of the LNBA lineage has not been explored, but a previous estimate of the species population size through time suggests it was comparable to extant *Y. pestis* ^19^. Nevertheless, it is possible that bottlenecks, which reduce the effective population size, contributed to the lack of any geographic differentiation of the LNBA lineage by repeated removal of genetic diversity decreasing the expected time to coalesce ^81^. However, repeated bottlenecks also are expected to lead to stochastic fixation of deleterious nonsynonymous mutations, weakening the strength of purifying selection ^82^, for which we cannot find evidence in the available genetic data. Ultimately, the influence of selection and demographic history is difficult to resolve with the data at hand. Thus, a complete explanation of the lack of geographic differentiation of the LNBA lineage is still wanting.

The temporal co-occurrence of the six earliest isolates indicates that the LNBA lineage was already established over a wide geographic range, which in turn suggests a highly efficient means of dispersal during this period. It is possible that serial reseedings could have occurred through a highly mobile host, such as migratory birds. Flyways play a major role in globally disseminating avian influenza today ^83^, and the known geographic distribution of the LNBA lineage closely matches that of the Black Sea-Mediterranean Flyway, the Asian–East African Flyway and the Central Asian Flyway, three overlapping bird migration routes that connect breeding grounds in Siberia with wintering grounds in Africa and the Indian subcontinent ^83^. If a Siberian breeding ground were to sustain a *Y. pestis* population, it could serve as the reservoir for annually disseminating a single lineage of plague over vast distances by hundreds of migratory bird species. It is known that other early forms of plague were circulating in northern Eurasia, as all ancient *Y. pestis* genomes basal to the LNBA lineage have so far been found in northern Europe ^6,50,84^. Extant *Y. pestis* has been identified in one bird species ^85^ and avian *Y. pseudotuberculosis* infections are prevalent ^86^. More than 1.5 billion birds are estimated to cross Central Asia every autumn ^87^, providing ample opportunity for direct or indirect disease transmission to local rodent or livestock populations. Why, however, extant *Y. pestis* no longer transmits over large distances, but instead is mostly seeded from local reservoir populations is difficult to answer but may be linked to its genomic changes.

Only by continuing to push paleomicrobiology from its human-centric beginning into the zooarchaeological record will we be able to better understand whether or not a given animal or transmission route was key in perpetuating plague in human populations during the LNBA. Because the underlying transmission cycle of the LNBA lineage was presumably multifactorial, including climatic, cultural, and economic parameters ^57,88^, multi-disciplinary research is needed to understand the emergence and spread of *Y. pestis* as well as other zoonotic pathogens during prehistory ^4,13–15,89^. Expanding the scope of current research to bring in new lines of evidence is needed to reveal the role of human-animal interactions in the evolution of human disease burdens.

## Acknowledgements

We thank the Key lab for helpful discussion, Diane Schad & Ronny Nitschke for graphical support. This work was supported by the Max Planck Society.

TH: This research was supported in part by the Wenner-Gren Foundation (No. 9973).

PK: We thank the Museum of the Institute of Plant and Animal Ecology UB RAS for providing samples for this research.

AH: Alexander Herbig was supported by the European Research Council (ERC) under the European Union’s Horizon 2020 research and innovation program (grant agreement no. 856453 ERC-2019-SyG)

CJ: Choongwon Jeong was supported by the National Research Foundation of Korea (NRF) grant funded by the Korea government (MSIT) (No. RS-2023-00212640) and by the Global-LAMP program of the National Research Foundation of Korea (NRF) grant funded by the Ministry of Education (No. RS-2023-00301976).

CW: American School of Prehistoric Research; Max Planck-Harvard Research Center for the Archaeoscience of the Ancient Mediterranean, European Research Council (StG-804884, DAIRYCULTURES); Max Planck Society; Werner Siemens Foundation (Paleobiotechnology); German Science Foundation (DFG EXC 2051 #390713860)

FMK: Felix M. Key is supported by the Klaus Tschira Foundation (GSO/KT030)

## Author Contributions

Conceptualization and Supervision: F.M.K. and C.W.

Investigation and Formal analysis: I.L.M., T.H., R.A.B., L.S., D.K., C.J.

Data Curation: I.L.M.

Resources: P.K., V.A.

Writing - Original Draft: I.L.M. and F.M.K.

Writing - Review & Editing: I.L.M., T.H., B.P.H., A.H., C.W., F.M.K.

## Supplemental information

**Document S1:** Figures S1-7

**Table S1**: Califf et al. region information plotted in Figure S5

**Table S2**: Genes carrying signatures of parallel evolution on external nonsynonymous mutations (N = 23) identified on the LNBA lineage

**Dataset S1**: Genetic screening data generated from faunal remains

**Dataset S2**: ARK017 *Y. pestis* capture sequencing and genome reconstruction mapping information

**Dataset S3:** Host PCA Nuclear reference data

**Dataset S4:** *Y. pestis* sample accessions and metadata

**Dataset S5**: Annotated mutation table for SNVs identified across all ingroup samples

**Dataset S6**: Screening MALT database genomes

**Dataset S7**: Host validation MALT database accessions

**Dataset S8**: *Y. pestis* and *Y. pseudotuberculosis* pangenome genomes

**Dataset S9**: Normalized breadth measurements in Figure 3C

## STAR Methods

### RESOURCE AVAILABILITY

#### Lead contact

Further information and requests for resources and reagents should be directed to and will be fulfilled by the lead contact, Felix M. Key.

#### Materials availability

This study did not generate unique reagents.

#### Data and code availability

- Raw sequencing data generated for this project have been deposited at the European Nucleotide Archive under the accession PRJEB78495. Metadata gathered from previous publications and used for the current analysis has been gathered and deposited in the github repository for this project https://github.com/ilight1542/ARK017_project
- Previously published data were reanalyzed for this work (**Dataset S4**).
- All original code used for the processing of genomic data and subsequent analysis can be found in the github repository for the project https://github.com/ilight1542/ARK017_project
- Any additional information required to reanalyze the data reported in this paper is available from the lead contact upon request.

### EXPERIMENTAL MODEL AND SUBJECT DETAILS

#### Permission statement

Permission to work on the archaeological samples was granted in 2021 by the curator of the Arkaim zooarchaeological collections at the Institute of Plant and Animal Ecology of the Ural Branch of the Russian Academy of Sciences (permit number 16353-2115/214), who is a co-author of the study (P.K.).

#### Archaeological information

Arkaim is a circular and fortified Bronze Age settlement with an outer diameter of 160 m enclosing over 67 rooms organized in a radial fashion ^90^. It is located in the Southern Urals region of Russia (Bredinsky District, Chelyabinsk Oblast; coordinates: 52.6492610, 59.5714430), and it was a major center of arsenical bronze metallurgy and early horse chariotry. It was part of the Sintashta-Petrovka cultural complex (ca. 2000–1600 BCE) that emerged in association with a rapid growth in bronze-producing pastoralist populations and heavy investment in defensive architecture, marking a major change from previous periods and suggesting a rise in shared material culture, social complexity, and violence ^70,91^. Competition for bronze and water resources and the protection of a growing network of long-distance trade routes may have fueled these conflicts, and it has been hypothesized that Sintashta-Petrovka social organization centered around military-equestrian aristocracies ^92^. Grave goods included a wide array of weaponry, especially relating to projectiles ^93^, which likely reflects a new style of combat involving javelins or arrows shot from lightweight spoke-wheeled chariots driven by horse teams. No evidence for agricultural production has been found at Arkaim or other Sintashta-Petrovka sites ^74^, but there is ample evidence of animal husbandry, with a focus on cattle, sheep, and horse pastoralism ^69^. We sampled 23 medium and large bovid skeletal remains from domestic contexts at Arkaim for genetic analysis.

### METHODS DETAILS

#### Radiocarbon Dating

The tooth sample ARK017 was radiocarbon dated at the Curt-Engelhorn-Zentrum Archäometrie (CEZA) in Mannheim, Germany. A sample of tooth root was prepared using a modified Longin method ^94^ with ultrafiltration to retain collagen >30 kDa. The sample (MAMS-68213) returned a ^14^C age of 3532 ± 19 BP, with 45.3% C, 5.4% collagen, a C:N of 3.2, and a δ^13^C of -21.4 ‰. The date was calibrated using OxCal v.4.4 ^43^ with the IntCal20 atmospheric curve ^44^, yielding a 95.4% calibrated date range of 1935-1772 cal BCE. All LNBA lineage and pre-LNBA lineage samples with previously published uncalibrated ^14^C dates were calibrated using the same IntCal20 curve, after applying any reservoir effect corrections provided in the source publication (RV2039, GLZ001, GLZ002), as noted in the code and metadata repository for this publication (see data availability statement).

#### Laboratory procedures for genetic screening data generation

Tooth sampling, genomic DNA extraction, and single-stranded DNA (ssDNA) library preparation were performed in a dedicated ancient DNA cleanroom facility at the MPI-EVA. Tooth sampling was performed using the archived protocol dx.doi.org/10.17504/protocols.io.bqebmtan. DNA extraction was performed using a modified version of the protocol ^95^, as described in dx.doi.org/10.17504/protocols.io.baksicwe. We used an automated protocol for producing ssDNA libraries ^96,97^. Library preparation included double-indexing by adding unique 8-mer index sequences at both P5 and P7 Illumina adapters. We performed shallow shotgun sequencing to determine host species and allow screening for pathogen DNA. Single-end 75 base pair (bp) sequences were generated for all shotgun libraries and sequenced on an Illumina HiSeq4000.

#### Pathogen screening and identification of *Y. pestis* in ARK017

Twenty three shallow shotgun datasets (**Dataset S1**) collected from Arkaim were screened for the presence of human and animal pathogens using nf-core/eager version 2.4.6 with metagenomic species identification and initial authentication carried out by MALT and post-processing using a custom MaltExtract and AMPS script from the HOPS software package ^33,98^. Briefly, raw sequencing reads were quality filtered and adapters were removed. Read pairs of paired end sequencing data were merged. Mapping was carried out first to the GRCh38.p13 human reference genome. Unmapped reads for each sample were converted to a fastq file and used as input for MALT with a custom microbial database (see **MALT screening database construction**). Postprocessing of the rma6 MALT output file was carried out by running AMPS from HOPS to create summary files and subsequently a custom post-processing R script to report whether any of the target pathogen species were present and exceeded certain heuristic thresholds. The customized post-processing script was allowed for adjustment of several parameters to the following filters: edit distance distributions ^33^ (0.8 on all reads, 0.7 on “ancient” reads with damage), minimum damage at terminal base (10% of eligible reads show damage), and minimum read distribution (‘readDis’) of 0.75 (proportion of unique bases aligned of all bases aligned to reference genome). Summary files output by the post-processing script were inspected to identify candidates for capture sequencing.

#### MALT screening database construction

A custom database was constructed for the screening pipeline that accommodates genetic representation of intra-species microbial diversity while minimizing the number of genomes per species for computational efficiency. All bacteria and archaea genomes available from RefSeq ^99^ at the assembly level of “Chromosome” and “Whole Genome’’ were considered, unless this removed the entire genera of interest, in which case a single contig or scaffold level genome was included, using a “Scaffold” level assembly when available. All species taxonomic IDs (taxids) with multiple genomes were clustered using MeshClust2 ^100^, which selected a representative genome from each cluster. Selected genomes were then validated as correctly labeled by running a blastn search of the first 10,000 bases on the nt database. Results from the accession of interest were masked. For each blast query, a list of taxids associated with the species taxid was generated, that included all subspecies and strains which had the same genus and species name but a different taxid. Genomes with at least three top bitscore hits (maximum value, may be shared across multiple blast query hits) matching a taxid within the generated list were included in the database.

In addition to the above genomes, all viral genomes and several eukaryotic genomes, including some pathogenic protists, were also included that would possibly have representation within the archeological samples. Conterminator ^101^ was used to identify eukaryotic reference sequences with putative contamination across kingdom domains using default parameters. Contigs shorter than 1000 bases or which had >5% contamination were discarded. The complete list of genomes included in the database, along with associated metadata is available in **Dataset S6**. The database was constructed on a 2TB node using the default malt-build command with step size 4 for use with MALT and MaltExtract ^33,102^.

#### In-solution capture enrichment for *Y. pestis*

After the identification of *Y. pestis* DNA sequences in ARK017, we performed three additional DNA extractions for this tooth and generated a total of 6 independent libraries that we enriched for *Y. pestis* DNA by performing an in-solution capture using a previously published capture probe set comprising the extant genetic diversity of *Y. pestis* as well as the *Y. pseudotuberculosis* outgroup (IP32953) ^28^. Three captured libraries were sequenced paired-end 75 bp on an Illumina HiSeq 4000 and three captured libraries were sequenced paired-end 100 bp on an Illumina NovaSeq X Plus. *Y. pestis* sequences were identified in all six captured libraries, and all capture sequencing of ARK017 were combined and validated by investigating the breadth of coverage across the reference genome, read stacking, plasmid coverage, edit distance distributions, and the presence of deamination patterns consistent with authentic ancient DNA data (Table 1, **Dataset S2**, **Figure S2**).

#### Host genus and species identification

Because a secure taxonomic assignment from incomplete remains for caprine domesticated sheep can be difficult to distinguish from wild sheep and goats, in addition to domesticated cattle and wild aurochs can be challenging, for the 23 skeletal remains in the study, we examined their DNA alignments to a database of 34 mitochondrial genomes of wild and domesticated ungulates with habitat ranges in central Eurasia (also including human and dog) using MALT ^36^ (**Dataset S7**).

After confirming ARK017 as belonging to the genus *Ovis*, to clarify the host species we projected genome wide SNV data from the ARK017 shotgun sequencing against a *O. aries* reference genome into the PCA space defined by SNP genotypes of domestic and wild *Ovis* species. For the ARK017 data, Illumina adapter sequences were trimmed and trimmed reads with ≧35 bp length were retained for mapping with AdapterRemoval v2.3.1 ^103^. The processed reads were aligned to the oviAri4 (accession: GCA_000298735.2) reference genome using bwa aln/samse v0.7.17 ^104^ with a relaxed edit distance parameter (-n 0.01) and seeding disabled (-l 9999). Duplicate reads were removed using DeDup v0.12.8 ^105^, and reads with mapping quality lower than 30 were filtered out using SAMtools v1.9 ^106^.

Genotypes were determined for 38.5 million single-nucleotide polymorphisms (SNPs) from the International Sheep Genomics Consortium (ISGC) SNP project, including 70 domesticated sheep, three North American bighorn sheep (*O. canadensis*), and two Dall sheep (*O. dalli*) from Bioproject ID PRJNA160933. Using PileupCaller (https://github.com/stschiff/sequenceTools), we randomly selected one allele per position (minimum mapping and base quality of 30), making the dataset individuals homozygous at each locus. The resulting files were merged with Plink v1.9 (Chang et al., 2015). To enhance the ISGC SNP panel, we incorporated published genome data of 16 Asiatic mouflons (*O. orientalis*). We obtained FastQ files for these individuals from the NCBI Sequence Read Archive (accession number PRJNA24020) and aligned the reads to the oviAri4 reference genome using BWA-mem v0.7.17 ^104^. PCR duplicates were removed with the MarkDuplicates module from picardtools v2.20.0, and properly paired reads with a mapping quality score of 30 or higher were retained using SAMtools v1.9 ^106^. For the ISGC SNP project SNPs, genotype likelihoods were calculated using GATK UnifiedGenotyper (v3.8.1.0) with the options ‘–genotype_likelihoods_model SNP –output_mode EMIT_ALL_SITES –allSitePLs’ ^107^. An in-house python script was then used to calculate posterior genotype probability with a non-default prior [0.4995, 0.0010, 0.4995] to minimize reference bias, as done in the Simons Genome Diversity Project. Genotypes with posterior probabilities of 0.9 or higher were retained, with the rest marked as missing. Modern wild and domestic sheep samples from the ISGC, 16 Asiatic mouflons are detailed in **Dataset S3**. PCA was conducted using smartPCA from EIGENSOFT with the option ‘lsqproject: YES’, projecting the ancient sequence from ARK017 onto the first three components defined by the full dataset (including wild sheep and domestic sheep references) to visualize diversity among sheep species ^108^.

#### Public *Y. pestis* genomes and raw sequencing data processing

Samples for reanalysis included all previously published LNBA lineage *Y. pestis* samples and representative samples from other ancient lineages ^6,8,19,27,28,84,109–118^. Additionally, samples representing the modern extant diversity were included by reprocessing all modern samples used in Valtuena et al. 2022 which had raw reads accessible ^8^. The final list of analyzed genomes can be found in **Dataset S4**. Raw sequencing data was downloaded using nf-core/fetchngs ^119^ and metadata regarding library preparation was retrieved from the ancientmetagenomicsDir Tools ^120^ Using the nf-core/eager software pipeline, ancient samples were pre-processed considering their individual library preparation, aligned to the *Y. pestis* reference CO92 (Accession number: ASM906v1) using bwa-aln (-n 0.01), deduplicated and, if non or half UDG treated had either 2 or 3 terminal bases of all aligned reads masked, respectively; modern sequencing efforts were processed as above, but bwa-mem was used for the alignment.

The resulting deduplicated bam files were merged across all UDG-treatments and libraries per sample and used for variant calling with a custom snakemake pipeline. Briefly, samtools mpileup ^106^ was used with minimum alignment mapping quality 30 to generate variant support across all samples at all positions. Then, bcftools was used to identify candidate SNVs which had a minimum allele frequency of 0.1, a loose setting which was used to collect information for downstream filtering steps. A total of 207,210 candidate SNVs were identified across all genomes. For each candidate SNV, summary statistics (base support counts, genotype likelihoods, coverage) across all samples were aggregated into multi-dimensional matrices utilized for subsequent quality control and analysis. Additionally, genome coverage bins were calculated for each contig within the reference genome using bedtools ‘genomecov’ and zero-coverage regions were identified using bedtools ‘genomecov -bgà, ‘mergè and ‘coveragè for downstream filtering of ancient datasets (see *Filtering variant sites for metagenomic misalignments) ^121^*. Information on the nf-core/eager runs, including all parameters used, the snakemake pipeline, and other data generation scripts is available in the project github repository (see data availability statement).

#### High quality variant identification and phylogenetic reconstruction

Output matrices from the snakemake pipeline and bedtools outputs were processed in a python (version 3.11.4) environment to carry out quality control and annotate variants following and extending upon a previously published toolkit ^122^. First, samples were excluded from downstream analysis if the average genome-wide coverage was below 2 (2 genomes removed).

##### Filtering variant sites within samples

For every candidate variant position identified by bcftools, we called a base in each sample by setting the highest supported base as the base call. We masked base calls on a sample-by-sample basis by setting calls to N if they were not covered by at least 2 reads with at least 1 read each mapping in the forward and reverse direction, if the FQ score (consensus quality produced by samtools) was greater than -30, if the major allele frequency (MAF) was below 0.9, or if more than 50% of reads supported an indel call at a position within 3 bases of the candidate variant. These metrics removed 200488 variants.

##### Filtering variant sites for metagenomic misalignments

Additional quality control metrics for bases were applied on a sample-by-sample basis for ancient genomes, reconstructed from metagenomic sequencing data of archeological remains. These samples contain genetic information not just from the target species, but also environmental species, which can result in erroneously alignment of orthologous sequencing reads and subsequently well supported but false variant calls. Here we apply two filter steps to mask base calls within a sample resulting from misalignments in ancient samples. First, to remove misalignments due to highly conserved regions across the species in the reference genome, we mask base calls in regions with coverage exceeding the 95^th^ percentile per contig analyzed (either chromosome or each plasmid). Second, isolated, short regions (< 500 bp) of coverage within otherwise non-covered genomic areas may be due to misalignments leading to spurious variant calls. We identified all regions with zero coverage, merged all these regions if they were within 500bp of each other and masked base calls in the region if the total proportion of this region with zero coverage exceeded 95 percent. Lastly, we masked variant calls if the base was within 10 bp of another candidate variant with a MAF < 0.9. These filters removed 674 variants present only on ancient samples.

##### Filtering variant sites across samples

We utilized information from across samples to identify regions with consistent issues in mapping. A variant position was masked if the covariance of the minor allele frequency across genomes was above the 95^th^ percentile (covariance is expected to be low for true variants not affected by mismappings), if more than 25% of retained samples had an uncalled or masked base on the position (limiting the influence of accessory genome content in the reference), if the median coverage across all genomes at the position was below 2 reads, or if the position had evidence of recombination (non-clock-like evolution), assessed by identifying SNVs less than 500bp apart whose allelic state covaried highly (above 99% percentile) across samples (assessed by calculating the covariances of the focal SNV with all other SNVs among ingroup samples). Lastly, a set of previously characterized repeat regions and RNA annotated regions in the CO92 reference genome were masked ^20^. These filters removed 1387 additional variants.

##### Filtering variant singletons in ancient samples

Finally, we further scrutinized singleton SNVs (positions with an alternative variant call supported in only one sample after above filtering), in ancient samples as these represent SNVs at risk to be contamination driven. For each ancient singleton, we extracted reads above Q30 supporting the position using samtools view and subjected each read to a blast query using the ‘-task blastn-short’ option against the nt database. Reads were discarded as contamination-driven if the top hit was not *Y. pestis* or *Y. pseudotuberculosis* based on bit-score and percent identity (considering all hits with the same maximum bit-score and percent identity). All relevant coverage-based matrices were updated and the variant was retained if the position still passed the MAF and coverage quality control thresholds. A total of 1003 singleton positions were investigated across ancient samples, of which 217 SNV positions were identified as stemming from contamination and discarded for downstream analysis.

##### Phylogenetic analysis

A maximum-likelihood (ML) phylogeny of all 200 retained samples and 4,444 retained variant positions was created using RAxML-ng version 1.2.2 using the GTR+G model and 1,000 bootstrap trees ^123^. The *Y. pseudotuberculosis* sample IP32953 was set as the outgroup. Felsenstein bootstrap proportions ^124^ were calculated and placed on the bestTreeCollapsed output from RAxML-ng using SumTrees within Dendropy ^125^. TreeTime was used to generate a maximum likelihood internal node reconstruction of the variant positions using the ‘ancestral’ module ^126^.

#### Gene content analysis

To identify the presence of any *Y. pseudotuberculosis* genetic content retained in the LNBA lineage of *Y. pestis* but not present in the CO92 reference genome, a separate mapping was carried out to the *Y. pseudotuberculosis* IP32953 reference genome (accession number: ASM4736v1). All data preprocessing and mapping was done with the nf-core/eager pipeline using the same parameters as for the mapping to *Y. pestis* CO92. Unfiltered, deduplicated bam files for each sample were merged across different UDG-treatments and bedtools was used to assess the average breadth and depth of coverage on each genic region in the *Y. pseudotuberculosis* IP32953 reference genome. Additionally, we mapped the sequences used for the capture-enrichment probe set, which were *in silico* designed including the *Y. pseudotuberculosis* IP32953 reference genome and utilized to reconstruct multiple human-derived *Y. pestis* genomes ^8,19,28,109–111,113,115,116,118^, in order to validate the mappability of sequences captured to the IP32953 reference genome.

We utilized the software packages Roary ^127^ and Prokka ^128^ to characterize the gene content of all “Complete” and “Whole-genome” level assemblies of *Y. pestis* (n = 66) and *Y. pseudotuberculosis* (n = 27) on NCBI RefSeq ^99^ (**Dataset S7**). First, Prokka was used to annotate the reference sequences using the ‘--genus Yersinià option. These annotated GFFs were then used as input by Roary to generate information about the presence of genic regions across the *Y. pestis* and *Y. pseudotuberculosis* assemblies. Using the output files of Roary, we defined a *Y. pestis*-*Y. pseudotuberculosis* “strict-core genome” as all genic regions present across all *Y. pestis* and *Y. pseudotuberculosis* reference genomes analyzed (2235 genes) and the *Y. pseudotuberculosis* “core genome” as all genic regions present in >90% of *Y. pseudotuberculosis* reference genomes analyzed (3081 genes). Using the output files of Roary, genes within the *Y. pseudotuberculosis* IP32953 genome feature file were mapped to the accessions names from the Roary output.

Prior to our characterization of *Y. pseudotuberculosis* core genes within the LNBA lineage and other branches of the *Y. pestis* diversity we applied two quality controls to the data. First, to control for biases due to variance in overall genome coverage, we normalized the breadth measurements across all genes by the average breadth of genome coverage on the *Y. pestis*-*Y. pseudotuberculosis* “strict-core genome” genes per sample. Breadth values of any gene which had normalized breadth values above 1 were set to 1. Second, since the only *Y. pseudotuberculosis* reference genome included for the *in silico* design of the *Y. pestis* in-solution capture probe set was IP32953, we subset the core genome breadth data to genes within that reference.

The PCA was conducted using scikitlearn version 1.3.0 from the normalized breadth data across *Y. pseudotuberculosis* core genes for all high-quality genomes retained during phylogenetic analysis. For barplots and p-value calculations of estimated gene content, we set a binary cutoff of gene presence at normalized breadth > 90% and summed within each sample. Since binary cutoffs are sensitive to variability of overall breadth measurements and small sets of observations are less robust to outlier samples, we removed outlier samples RT6, ARK017, C10098 and Gok2, whose sum of gene presence (normalized breadth > 90%) across strict-core genes was more than two standard deviations below that of all analyzed samples, since the strict-core genes are expected to be present in all *Y. pestis* and *Y. pseudotuberculosis* genomes. Then, we compared the LNBA lineage genomes to extant genomes, early ancient genomes recovered from time periods contemporaneous or prior to the LNBA lineage genomes (RV2039, RT5, I2470), and late ancient genomes (1st/2nd Pandemic genomes) using a Mann-Whitney-U test. Barplots display the mean of each set of samples, with a 95% confidence interval. See **Dataset S4** for information on sample groups. We calculated the correlation of sample age and loss of total core gene content by pairing observations of summed breadth scores across core genes and calibrated BCE ^14^C dates of LNBA lineage samples, using a linear regression and sample median age as the predictor variable.

To visualize the gene content present in the LNBA lineage or basal lineages but lost in the diversity of extant *Y. pestis*, we plotted normalized breadth values across all samples in the phylogenetic tree for genes for which at least one LNBA lineage or basal ancient sample (Gok2, RV2039) had a >99% normalized breadth value (inferred gene presence) and more than 75% of samples belonging to Branch 0 to 4 had a <90% normalized breadth value (inferred gene absence or partial deletion). For visual clarity, we collapsed the breadth value across each branch 0 lineage, Branches 1-4, RT5/RT6/I2470, and 1st/2nd Pandemic genomes, respectively, displaying the maximum breadth value observed in each group. Normalized breadth measurements across all samples for these genes are shown in **Dataset S8**. Lastly, we explicitly investigated genic regions in *Y. pseudotuberculosis* IP32953 that have previously been characterized as being predominantly lost in *Y. pestis* (**Table S1**, **Figure S5**) ^46^. All heatmaps are ordered by the genic location and phylogenetic pattern of samples. A simplified phylogeny was matched to the samples displayed according to their order of the tree.

#### Identification of non-neutral evolution

We characterized genic SNVs as nonsynonymous if the variant caused an amino acid change or premature stop codon relative to the ancestral state of the position. We defined the ancestral state as the maximum likelihood state in the most recent common ancestor node (MRCA) using the software package TreeTime’s ancestral reconstruction output ^126^. To directly compare the extant diversity and the LNBA lineage, the MRCA of these samples was used as the ancestral state.

We generated SNV sets arising in different sections of the *Y. pestis* diversity to investigate possible signals of non-neutral evolution (see **Dataset S5**). We identified all SNVs arising on the LNBA lineage and the extant diversity by assessing if LNBA lineage samples or extant samples had a mutation relative to the MRCA of the LNBA lineage and the extant diversity. We also generated a set of SNVs arising on external nodes of the LNBA lineage, the extant diversity, and the human infection samples from the extant diversity.

We calculated dN/dS for the different sets of SNVs by normalizing the observed N(onsynonymous)/S(ynonymous) ratio by the expected N/S ratio. The expected N/S ratio was calculated by considering the genome-wide codon composition as well as the mutation frequency spectrum identified from all high quality SNVs. The 95% confidence interval is calculated using a Wilson score interval, implemented in the python package statsmodels version 0.13.5.

Parallel evolution, multiple independent SNVs occurring within the same gene, was quantified on two levels. First, whether or not the total number of genes observed with multiple mutations was unexpected when compared to 10,000 simulations of the same amount of randomly distributed genic mutations. Secondly, a p-value was calculated for whether the number of SNVs falling within a specific gene was significantly different from expectations under a Poisson distribution by using a survival function based on the per-base mutational probability given the number of mutations in the set, the length of the gene in question and the number of mutations in that gene. P-values were corrected for multiple testing using the Benjamini-Hochberg FDR correction at a significance level of 0.05, implemented in statsmodels.

### KEY RESOURCE TABLE

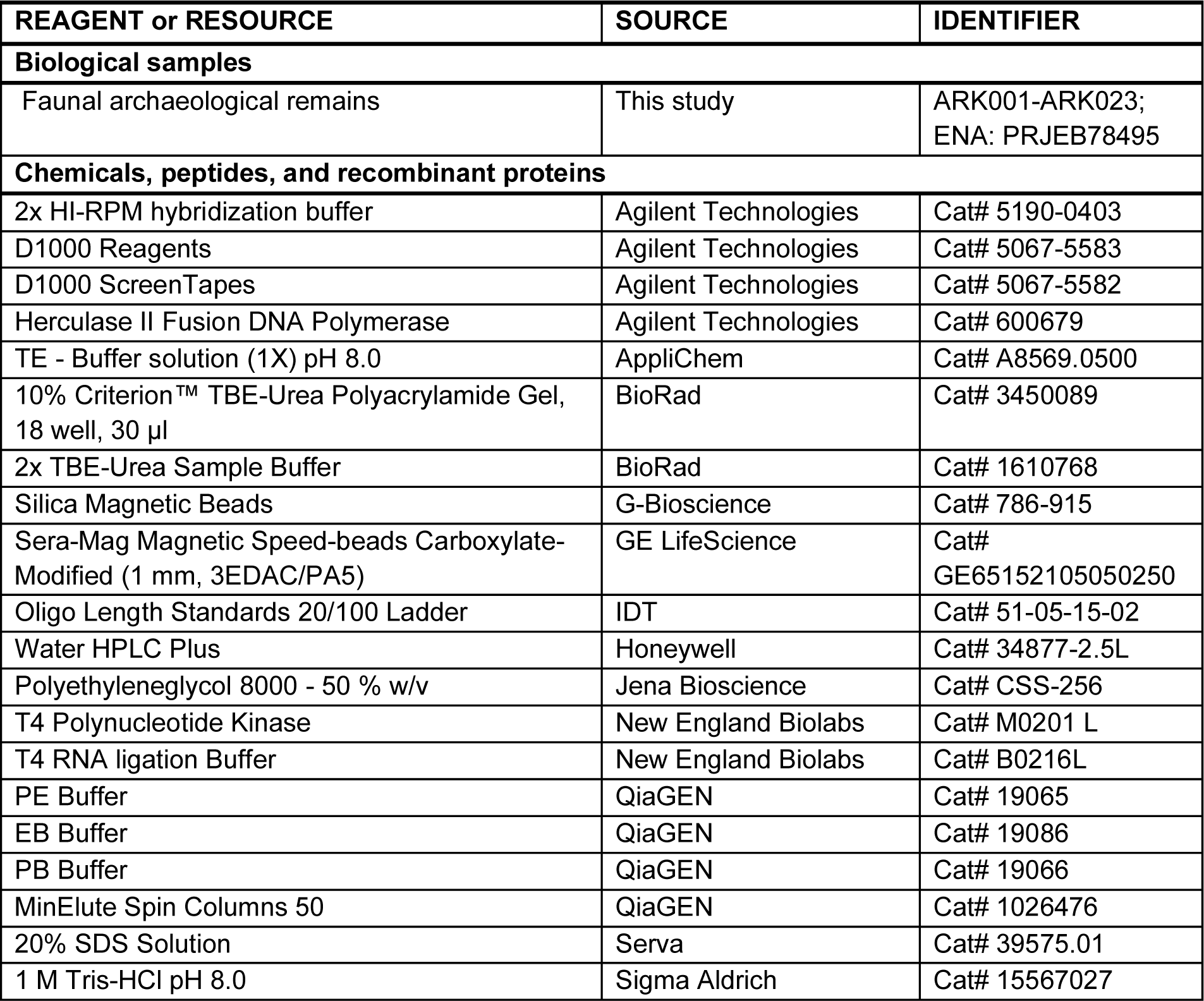
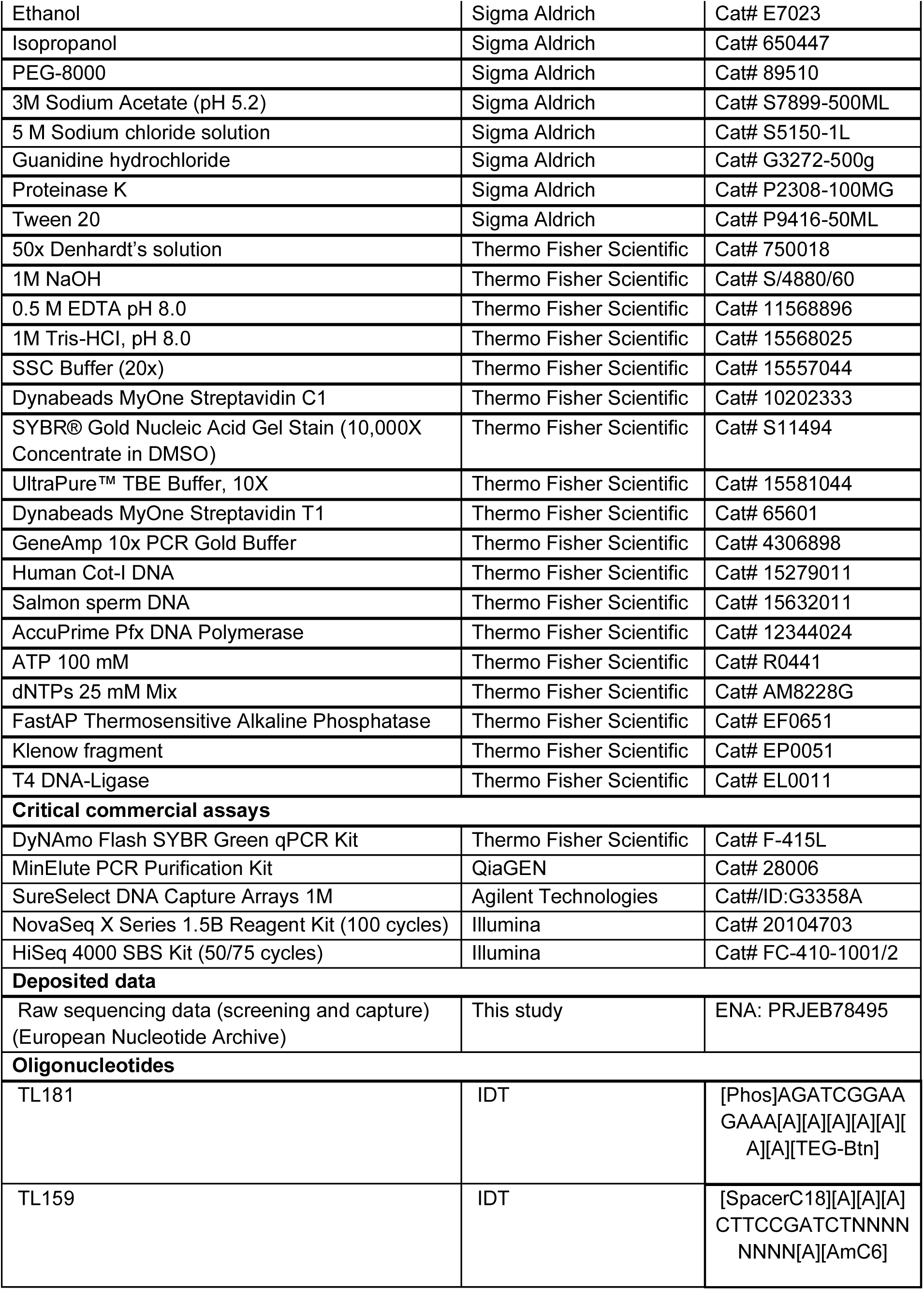
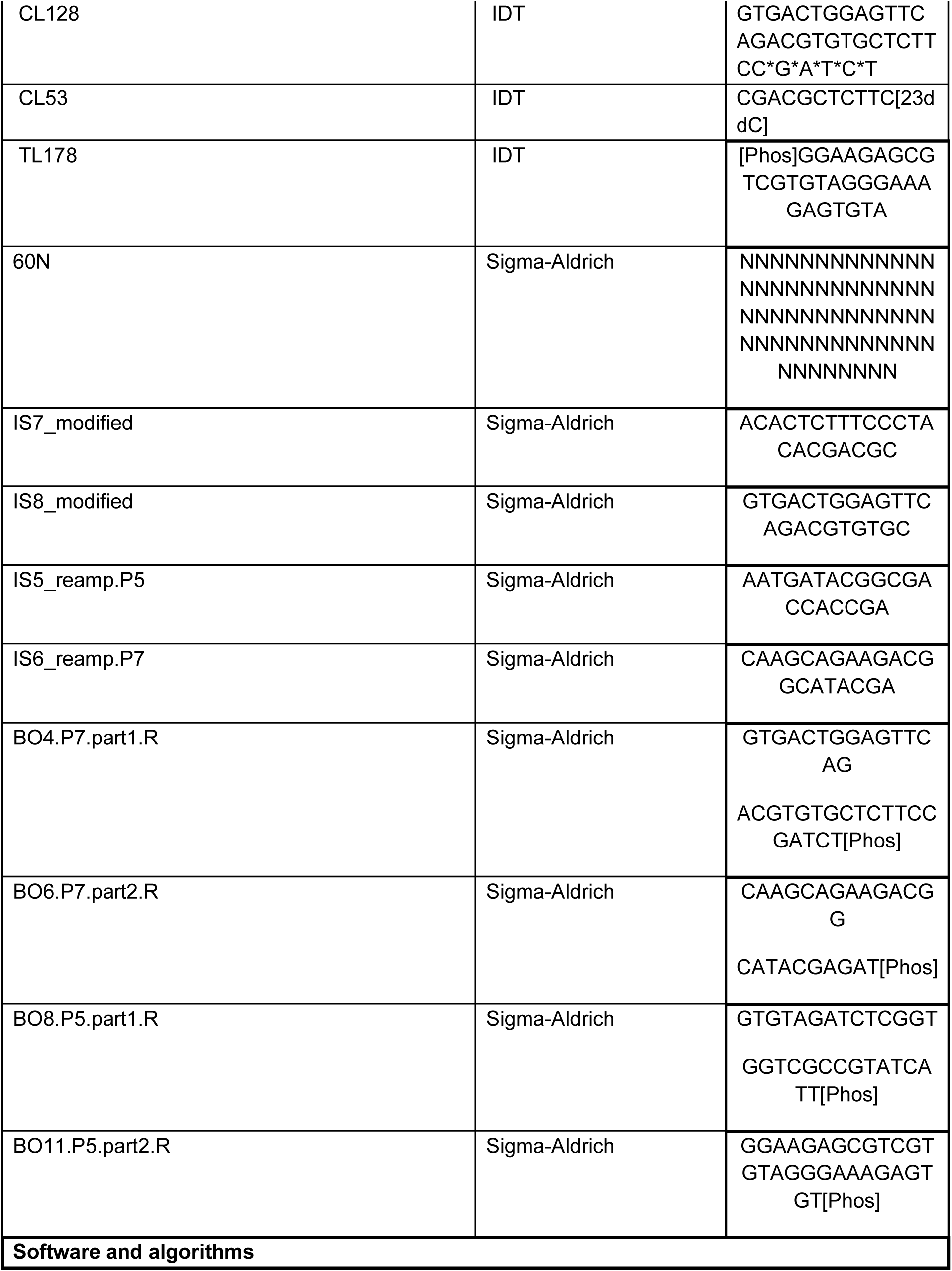
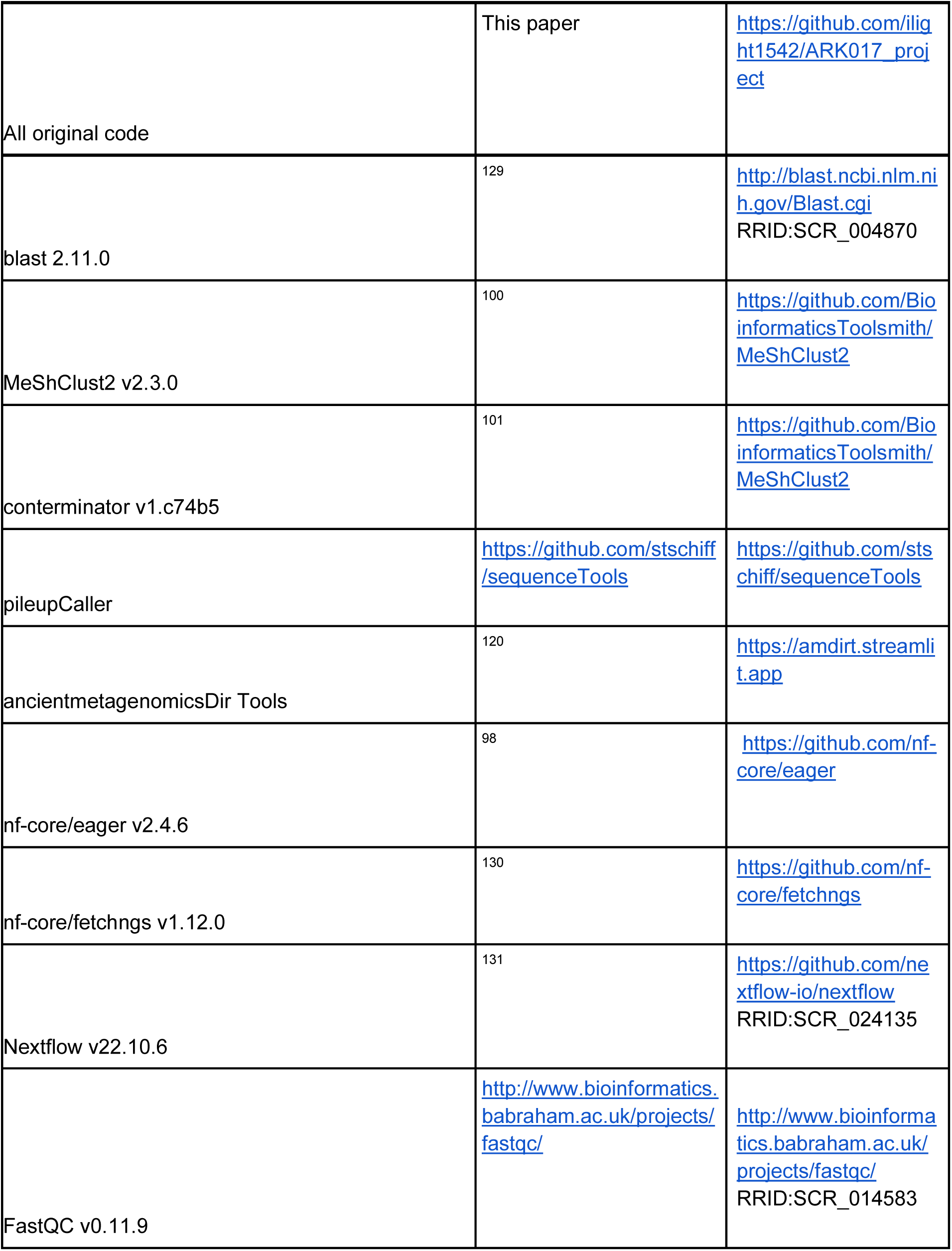
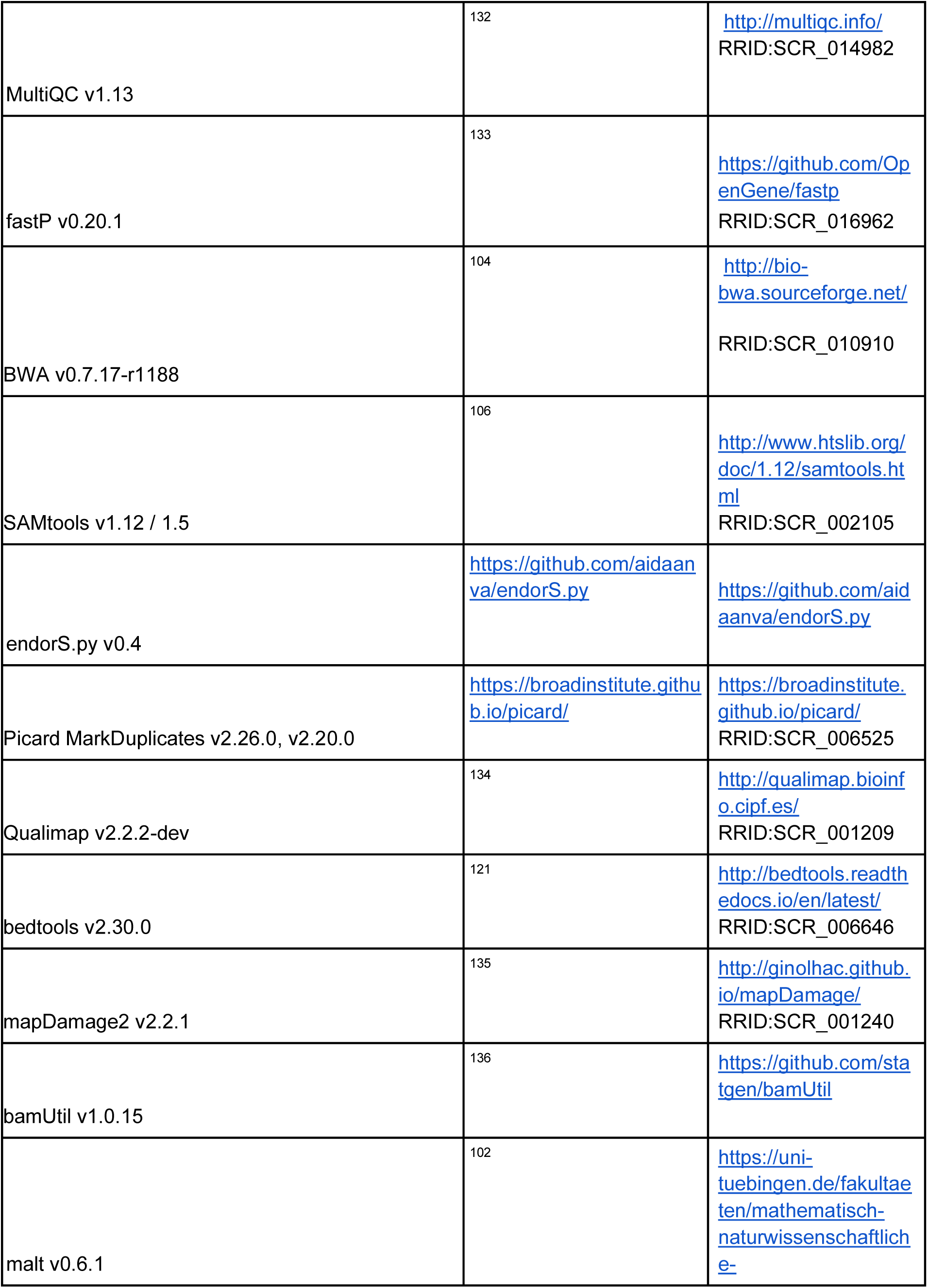
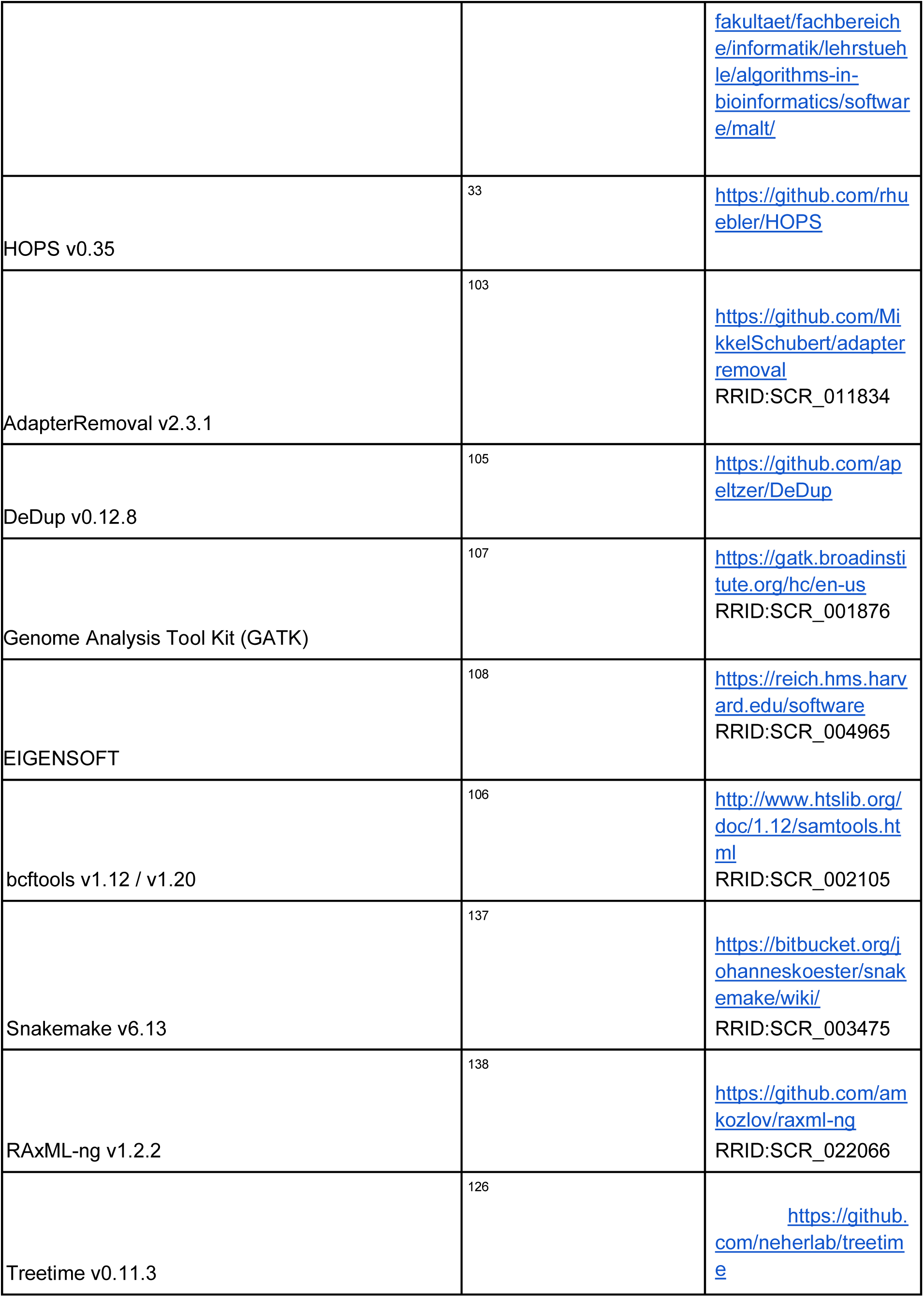
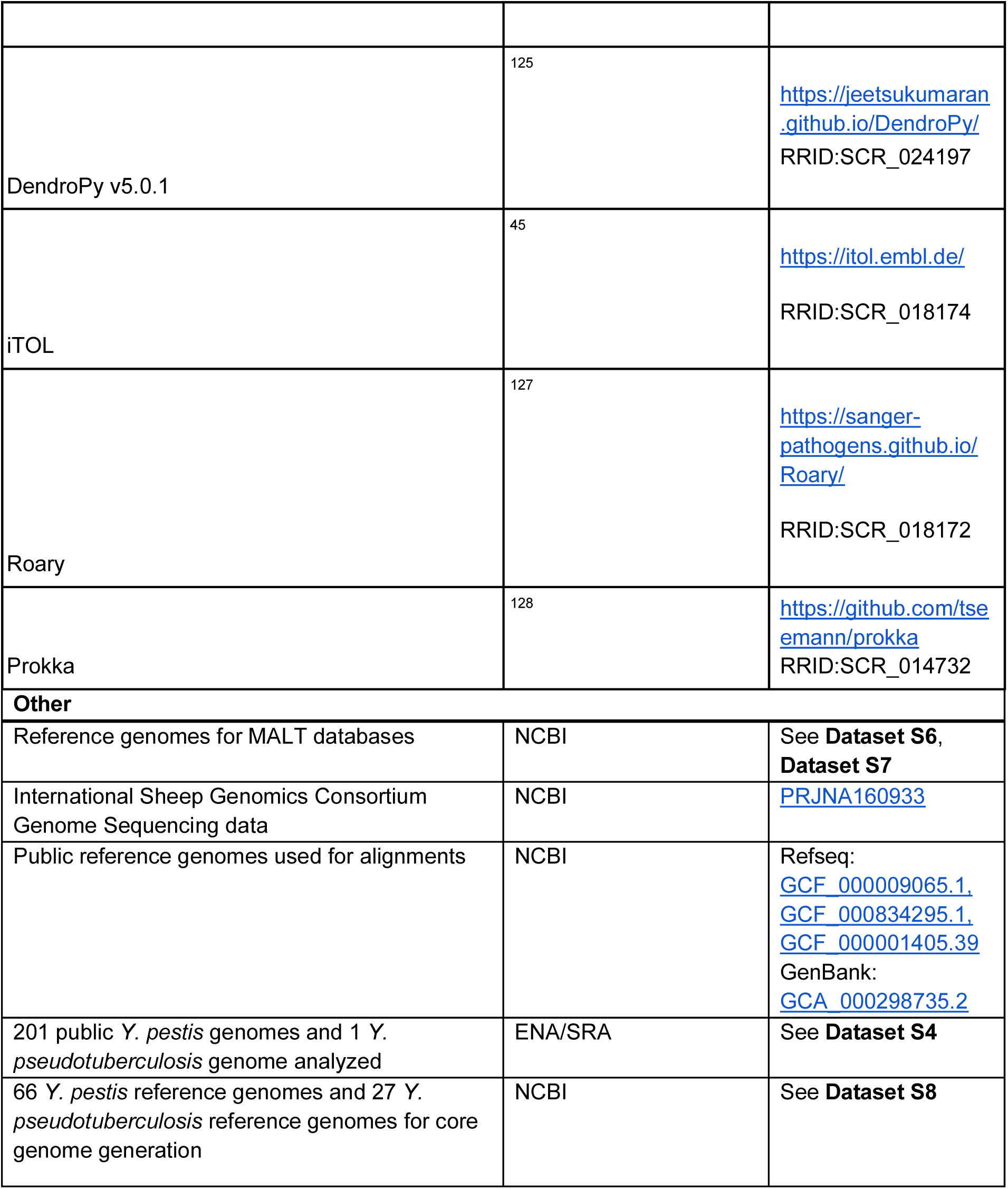

