## Supplementary Figures 1-7 for "Bronze Age *Yersinia pestis* genome from sheep sheds light on hosts and evolution of a prehistoric plague lineage"

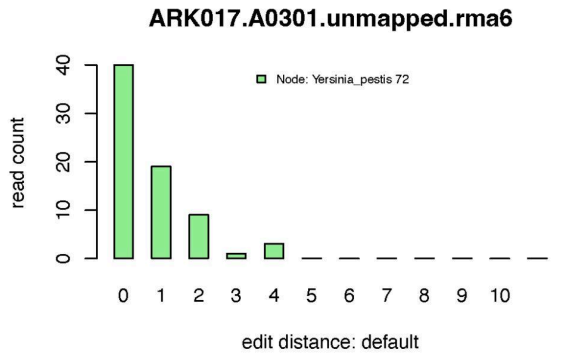


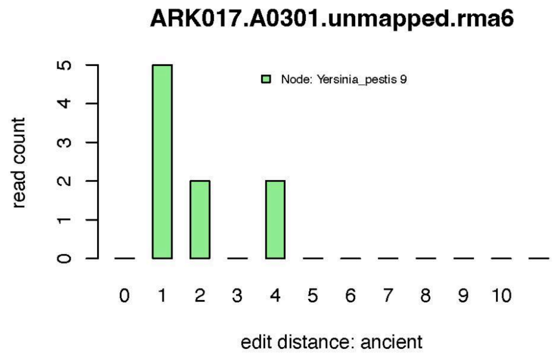


| | |
|---|---|
| Node | Yersinia_pestis |
| Top Reference | Yersinia_pestis |
| all reads | 64 |
| nonDup | 44 |
| readDis | 0.941 |
| nonStacked | 42 |
| destacking? | Off |
| C>T_1 | 0.2021 |
| G>A_-1 | 0 |
| mean length (sd) | 46 (9.345) |

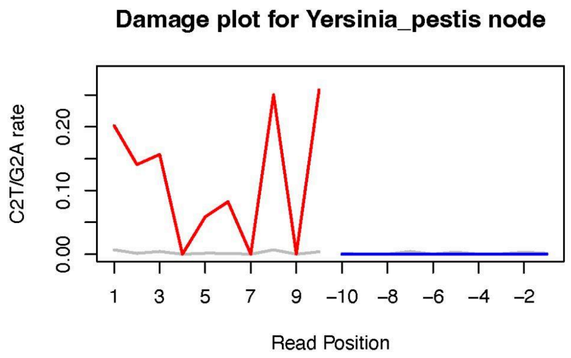


| |
|---|
| Yersinia_pestis;_NZ_CP064125.2 100% |
| Yersinia_pestis_EV_NIIEG;_NZ_CP065918.1 100% |
| Yersinia_pestis_Pestoides_F;_NC_009381.1 100% |
| Yersinia_pestis_subsp._pestis_bv._Medievalis;_NZ_CP045158.1 100% |
| Yersinia_pestis_3067;_NZ_CP006754.1 100% |
| Yersinia_pestis_Antiqua;_NZ_CP009906.1 100% |
| Yersinia_pestis_1412;_NZ_CP006783.1 100% |
| Yersinia_pestis_8787;_NZ_CP006748.1 100% |
| Yersinia_pseudotuberculosis;_NZ_CP009757.1 100% |
| Yersinia_pseudotuberculosis;_NZ_CP033715.1 100% |

**Figure S1. Initial identification and authentication of *Y. pestis* DNA in ARK017 from shotgun sequencing data.** The software tool HOPS was used to screen the dataset and identify potential pathogen DNA signatures. The results for the taxonomic node *Yersinia pestis* are shown. The declining edit distances on all reads (top left) and reads with deamination patterns (top right) consistent with ancient damage are highly indicative of a true positive, as are the number of non duplicate reads, which are largely non-overlapping (‘readDis=0.94’ shows proportion of unique bases aligned of all bases aligned to reference genome) and the presence of 5’ end C-to-T transitions on 20.2% of reads indicative for ancient DNA damage. No G-to-A transitions are expected on the 3’ end of the sequencing molecule since a single-stranded library preparation was used.

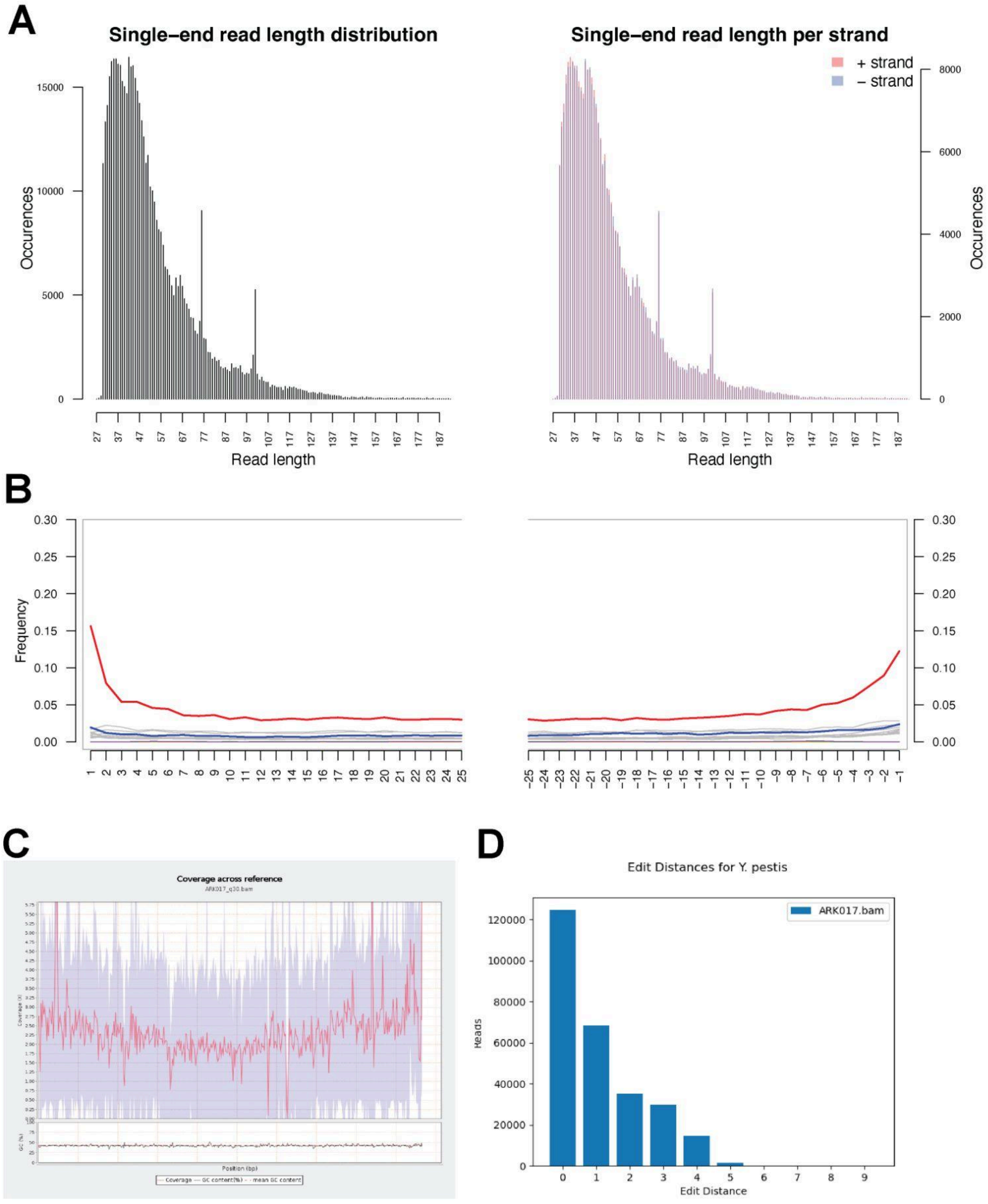


**Figure S2. Authentication of ancient origin of reconstructed *Y. pestis* genome ARK017 after in-solution *Y. pestis* capture. A.** ARK017 capture-sequenced reads aligned to *Y. pestis* CO92 reference are predominantly short fragments, consistent with ancient origin. Data shown is all capture-sequenced data merged after mapping and deduplication. Plot generated with mapDamage version 2.2.1 and the `--single-stranded` option. **B.** ARK017 capture-sequenced reads aligned to *Y. pestis* CO92 reference show deamination patterns consistent with ancient origin and single-stranded library preparation. Red line displays rates of C->T transitions on aligned reads for given locations on the read. Blue line displays rates of G->A transitions. Other transitions shown in gray. X-axis displays position on read; positive values are distance from 5' end and negative values are distance from 3' end. Plot generated by mapDamage, as in A. **C.** ARK017 capture-sequenced reads show even coverage across the reference genome. Data shown is all capture-sequenced data merged after mapping, deduplication, masking of 3 bases at the start and end of the aligned reads and subsetting to reads with at least Q30 in mapping quality. Plot generated with QualiMap version 2.2.2-dev using default settings. **D.** Edit distance distribution of all *Y. pestis* aligned reads from ARK017 sequencing after masking the first and last two bases of every read. See also **Dataset S2** for information about sequencing results of individual libraries.

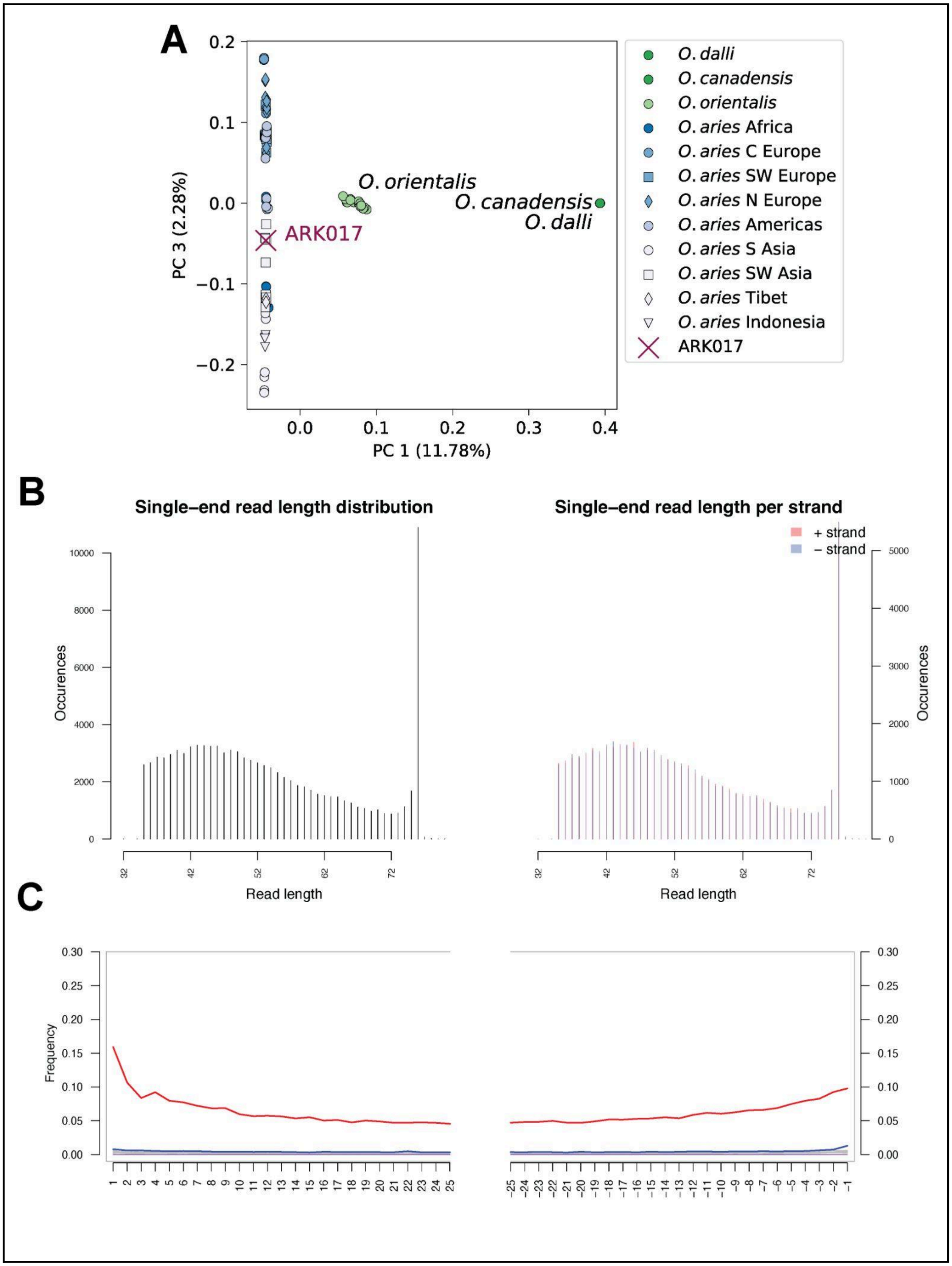
A
PC 3 (2.28%)
PC 1 (11.78%)
O. orientalis
O. canadensis
O. dalli
ARK017
O. dalli
O. canadensis
O. orientalis
O. aries Africa
O. aries C Europe
O. aries SW Europe
O. aries N Europe
O. aries Americas
O. aries S Asia
O. aries SW Asia
O. aries Tibet
O. aries Indonesia
ARK017
B
Single–end read length distribution
Single–end read length per strand
+ strand
– strand
Occurences
Read length
C
Frequency

**Figure S3. Authentication of *Ovis aries* host from shotgun dataset ARK017 A.** ARK017 *Ovis* ntDNA clusters with *Ovis aries* species from SW Asia. PC 1 separates Arkaim (ARK017) from other *Ovis* species, and PC 3 separates diversity among different modern *Ovis aries* populations using SNPs from the International Sheep Genomics Consortium SNP project. ARK017 clusters with the SW Asian representatives of the domesticated sheep *O. aries*. **B.** ARK017.A0301 shotgun-sequenced reads aligned to the *Ovis aries* reference genome (oviAri4) are predominantly short fragments, consistent with ancient origin. Data shown is shotgun-sequenced ARK017.A0301 screening data after mapping and deduplication. Plot generated with mapDamage using the `--single-stranded` option. **C.** ARK017.A0301 shotgun-sequenced reads aligned to the *Ovis* show deamination patterns consistent with ancient origin and single-stranded library preparation. Red line displays rates of C->T transitions on aligned reads for given locations on the read. Blue line displays rates of G->A transitions. Other transitions shown in gray. X-axis displays position on read; positive values are distance from 5' end and negative values are distance from 3' end. Plot generated with mapDamage, as above in C.

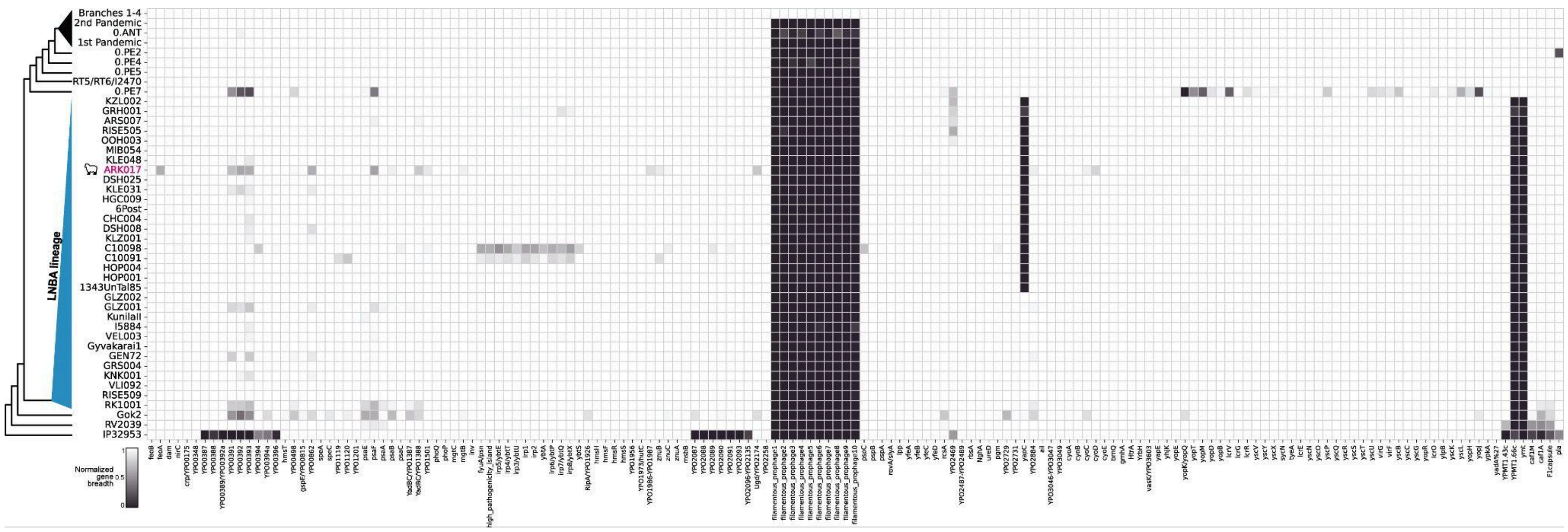

**Figure S4. CO92-based virulence gene presence and absence patterns.** Normalized gene breadth values were generated as in the main text except the values were normalized across all virulence genes analyzed in Andrades Valtueña et al. 2022. The results observed are in line with those previously reported and ARK017 presence/absence patterns agree with its placement within the LNBA lineage phylogeny. Collapsed values are displayed as described in **Figure 3A,** see also **Dataset S4**.

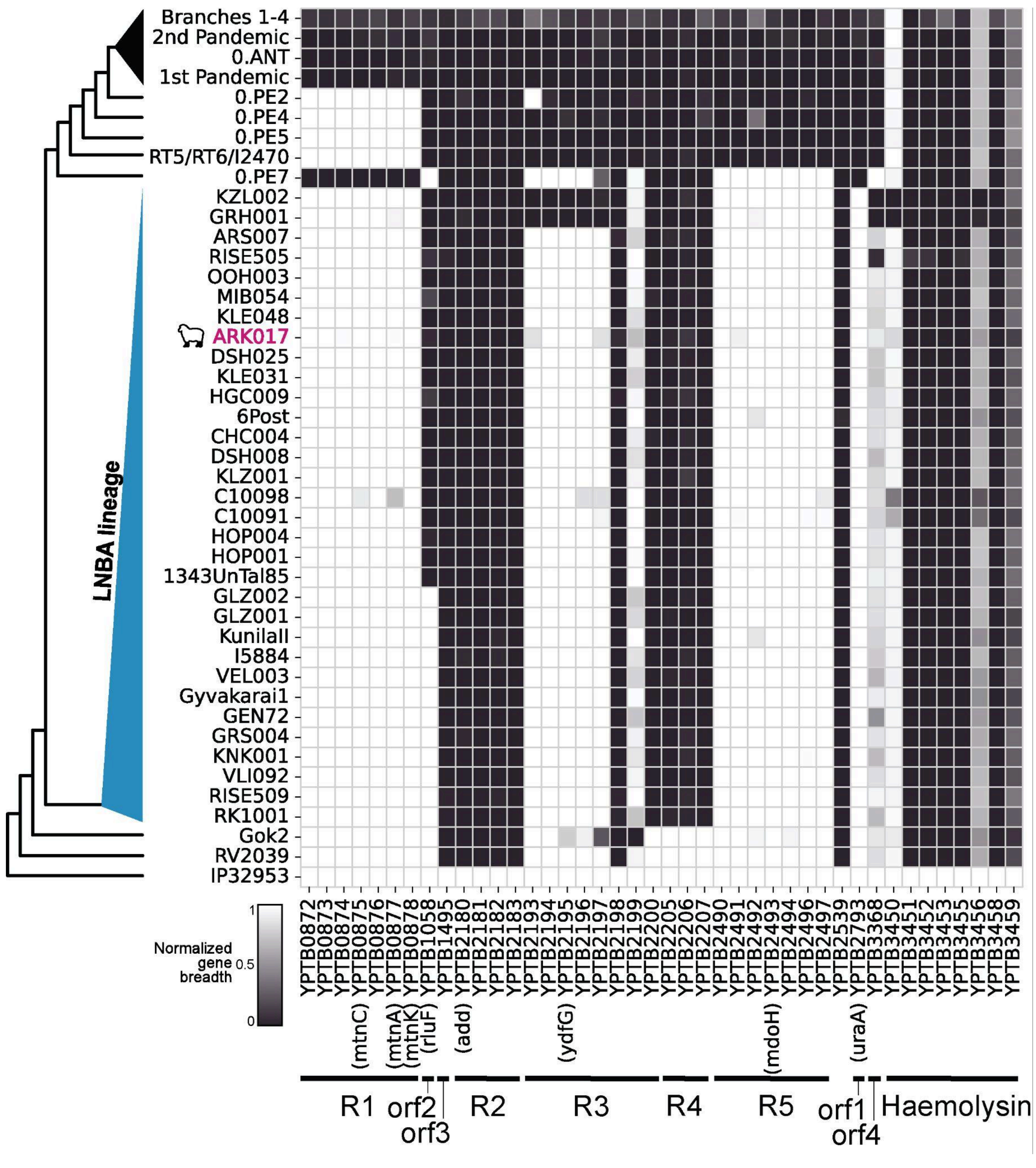


**Figure S5. Predominantly *Y. pseudotuberculosis* regions identified by Califf et al 2015** (Califf et al. 2015)**.** Normalized breadth measurements across regions investigated in Califf et al. 2015 and identified as mostly absent in *Y. pestis* but present in *Y. pseudotuberculosis*. Regions plotted even if they were not annotated as part of the *Y. pseudotuberculosis* core genome. Collapsed values are displayed as described in **Figure 3a,** see also **Dataset S4**, **Table S1**.

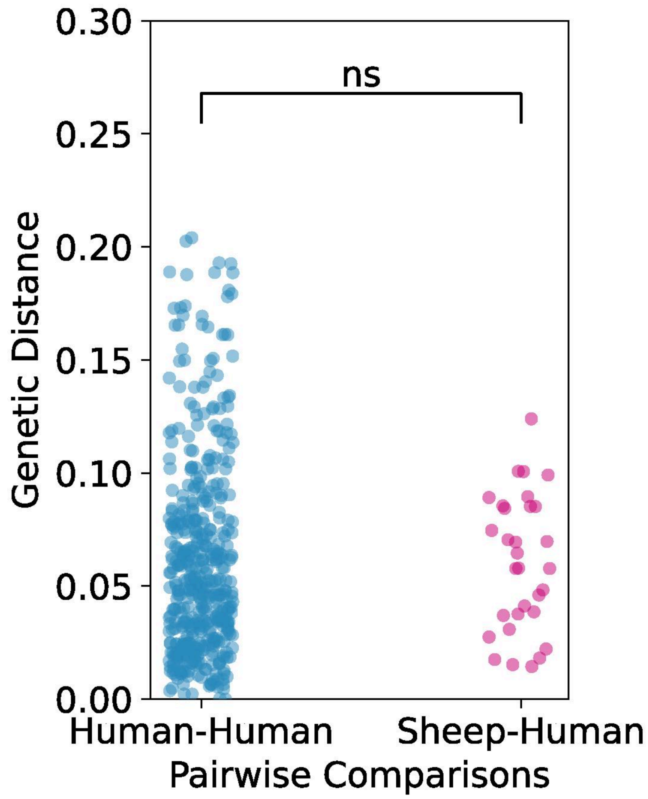


**Figure S6. Human-Human pairwise divergences (Non-ARK017) and ARK017-human pairwise divergences.** All pairwise comparisons of genetic distance for human-human comparisons and ARK017-human comparisons on the LNBA lineage. P-value=0.454 (Mann-Whitney-U test) shows the ARK017 genome is interleaved with human-derived, contemporaneous genomes. See also **Figure 4A**.

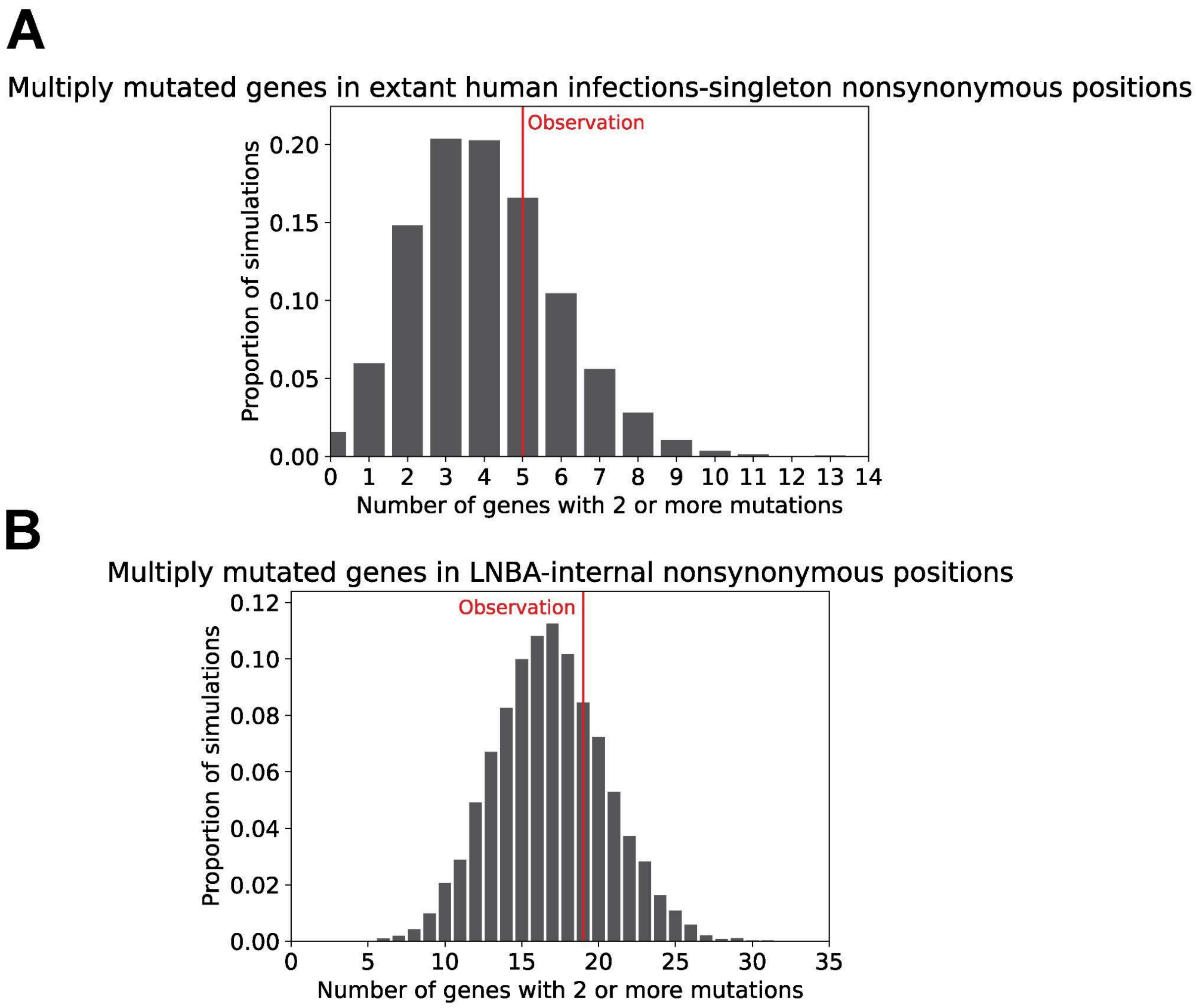


**Figure S7. Parallel evolution simulations across nonsynonymous positions mutation sets A.** Extant lineage human infections external nonsynonymous mutations mutate 5 genes multiple times (red). Simulated distribution of the number genes with multiple mutations (≥ 2) given the number of observed mutations (149 nonsynonymous mutations). **B.** LNBA lineage internal nonsynonymous mutations mutate 19 genes multiple times (red). Simulated distribution of the number of genes with multiple mutations (≥ 2) given the observed number of mutations (316 nonsynonymous mutations).
